# A Modular Platform for the Optogenetic Control of Small GTPase Activity in Living Cells Reveals Long-Range RhoA Signaling

**DOI:** 10.1101/2025.09.07.674731

**Authors:** Benjamin Faulkner, Yuchen He, Daniel Sitrin, Linda Ziamanesh, Cliff I. Stains

## Abstract

Small GTPases are critical regulators of cellular processes, such as cell migration, and comprise a family of over 167 proteins in the human genome. Importantly, the location-dependent regulation of small GTPase activity is integral to coordinating cellular signaling. Currently, there are no generalizable methods for directly controlling the activity of these signaling enzymes with subcellular precision. To address this issue, we introduce a modular, optogenetic platform for the spatial control of small GTPase activity within living cells, termed spLIT-small GTPases. This platform enabled spatially precise control of cytoskeletal dynamics such as filopodia formation (spLIT-Cdc42) and directed cell migration (spLIT-Rac1). Furthermore, a spLIT-RhoA system uncovered previously unreported long-range RhoA signaling in HeLa cells, resulting in bipolar membrane retraction. These results establish spLIT-small GTPases as a versatile platform for the direct, spatial control of small GTPase signaling and demonstrate the ability to uncover spatially defined aspects of small GTPase signaling.

## Introduction

Small GTPases are a diverse family of protein switches in mammalian cells and play pivotal roles in regulating a variety of essential cellular processes including cell survival, cell morphology, vesicle trafficking, and cell movement.^1–4^ In normal physiology, the activity of small GTPases is regulated via cycling between a guanosine triphosphate (GTP)-bound active state and a guanosine diphosphate (GDP)-bound inactive state (**Figure 1a**).^5^ This cycle of activation and inactivation is mainly regulated by two regulatory proteins: guanine nucleotide exchange factors (GEFs), which facilitate the exchange of GDP for GTP, and GTPase-activating proteins (GAPs), which accelerate the hydrolysis of GTP to GDP.^6^ Furthermore, a lipidation sequence present at either the N- or C-terminus of small GTPases can influence localization within a cell.^7,8^ Given their essential roles in cellular processes, mutations that perturb small GTPase regulation contribute to a range of diseases including cancer,^9,10^ neurodegenerative diseases,^11–13^ cardiovascular diseases,^14–16^ and immune disorders.^17–19^ Additionally, the redistribution of small GTPase activity within a cell upon treatment with chemotherapeutics has been shown to confer drug resistance.^20,21^ As such, understanding the molecular mechanisms that govern small GTPase activity at the subcellular level and developing tools to modulate their function with spatial precision remains a critical goal in biomedical research.

**Figure 1.**
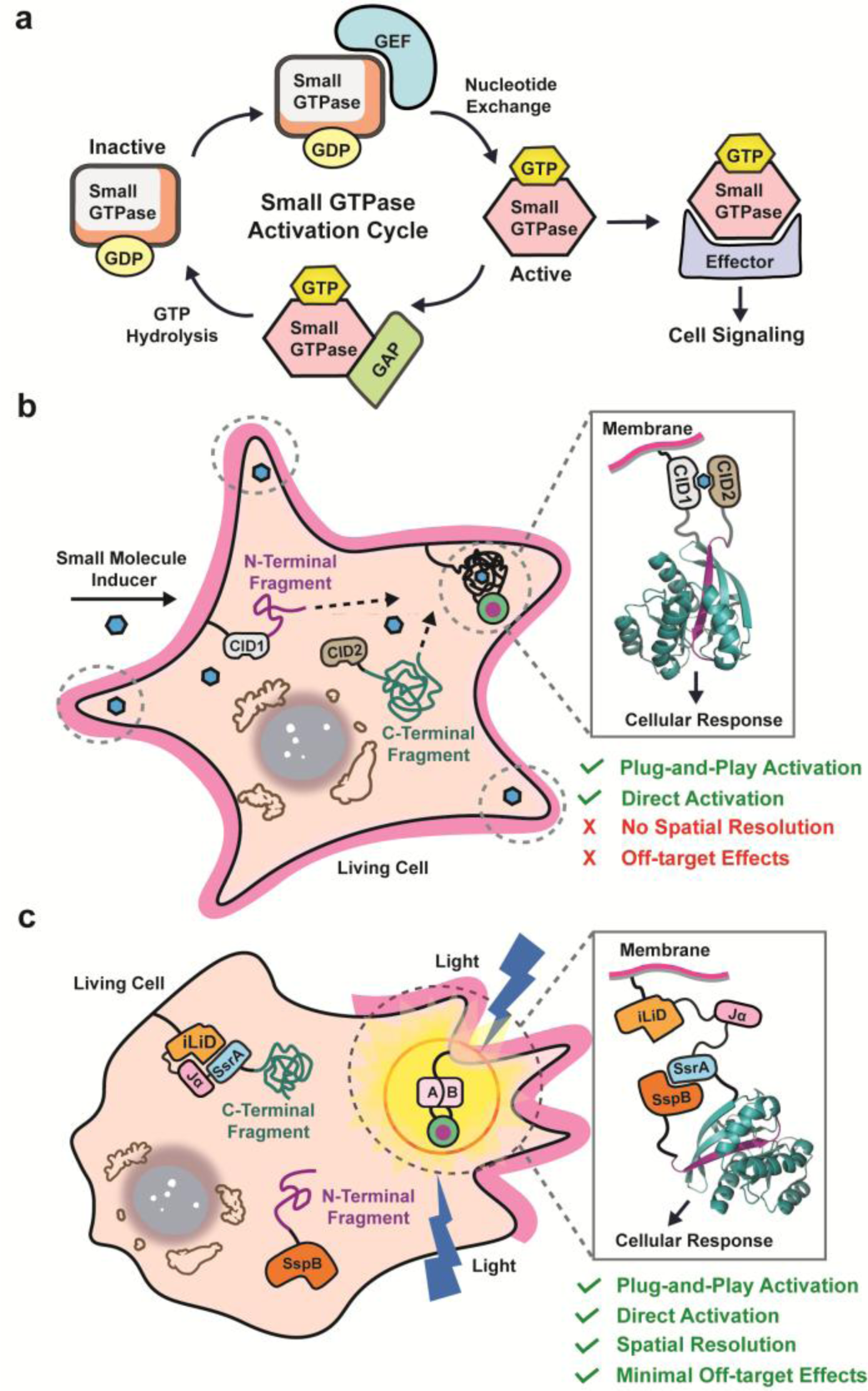
Split-protein engineering approaches to control small GTPase activity. **a)** Schematic illustrating the regulatory cycle of small GTPase activity. **b)** Fusion of small GTPase fragments to CID domains allows for the conditional reassembly and activation of small GTPases with user-defined inputs. **c)** Utilizing optogenetic domains (e.g. iLID) to control the reassembly of small GTPase fragments allows for spatial resolution of split-small GTPase activation.

Since the discovery of the Rat sarcoma (Ras) oncogene in the early 1980s,^22^ efforts to develop small molecule inhibitors to regulate small GTPase activity have received intense interest. However, these efforts have faced significant challenges due to the strong affinity of small GTPases for their natural substrates and an apparent lack of accessible allosteric binding pockets.^23^ Progress has been made recently in targeting small GTPases through covalent modification of residues near the GTP-binding site, a strategy that has since led to the first generation of small GTPase inhibitors entering the clinic.^24–29^ Nonetheless, this approach lacks the spatial resolution required to dissect the influence of subcellularly localized small GTPase activity. As a result, protein engineering-based approaches have gained increased attention for their ability to offer more precise, spatiotemporally controlled activation of small GTPase activity.^30–34^

Hahn and colleagues demonstrated the ability to control the activity of the small GTPase Rac1 by fusing a light, oxygen, or voltage (LOV) domain to the N-terminus of a constitutively active Rac1 mutant, blocking effector binding and rendering it inactive until exposure to blue light (450 nm – 500 nm).^35^ This light-activated system offers high temporal and spatial resolution for controlling small GTPase activity in living cells. Although, its reliance on a noncovalent binding interface between the LOV domain and Rac1 poses challenges for broad application across the small GTPase family, requiring intensive case-by-case optimization for each small GTPase. For example, when used to control the activity of the closely related small GTPase Cdc42, mutations had to be made in Cdc42 to obtain proper occlusion of its effector binding site in the dark state.^35^ Development of an alternative approach utilizing protein sequestration to localize GEFs to the inner cell membrane, known as LOVTRAP, allowed for a more generalizable approach to control the activity of other membrane-localized small GTPases such as RhoA and Rac1.^36^ Subsequently, Toettcher and colleagues developed the OptoSOS system to spatially regulate the activity of a RasGEF, SOS, via the selective localization of the GEF to the cell membrane using a light-responsive dimerization system.^37,38^ This method proved effective to study ERK signaling dynamics involved in cell proliferation.^38,39^ Although powerful, the cross-reactivity of GEFs with multiple small GTPases can complicate analysis of individual small GTPase outputs within complex cellular circuits. Thus, there is no general method for the direct, spatial activation of small GTPase signaling in living cells.

We have previously described a standardized parts set for the plug-and-play activation of small GTPases using chemical-inducible dimerization (CID) domains.^34,40,41^ By leveraging the high sequence and structural homology among small GTPases, we applied a fragmentation site originally identified in Cdc42^42^ to other small GTPases across the superfamily, including Rac1, RhoA, and KRas, without the need for case-by-case optimization (**Figure 1b**). Fusion of the relevant split-small GTPase fragments to CID domains enabled the small molecule-driven reassembly of each small GTPase, allowing for temporal control of small GTPase activity in living cells. While this approach allows for the potential dissection of small GTPase signaling within living cells, certain applications can suffer from the lack of spatial resolution and off-target effects arising from the use of CIDs to control split-small GTPase reassembly. Herein, we sought to create the first generalized approach for controlling the subcellular activation of small GTPase signaling through optogenetically-controlled reassembly of split-small GTPases (**Figure 1c**). The resulting platform enables localized activation of small GTPase signaling with high spatiotemporal resolution and can be paired with the appropriate split-small GTPase to modulate a chosen cellular phenotype. Additionally, we demonstrate that the extent of signaling activation in this system can be rationally tuned. Utilizing this platform, uncovered previously unreported bipolar membrane retraction in HeLa cells following localized RhoA activation, revealing long-range propagation of RhoA signaling across the cell. In the long term, we envision that our approach will enable the spatial dissection of cellular signaling pathways in biomedical research as well as applications in synthetic biology.

## Results and Discussion

### Light-Gated Activation of spLIT-Cdc42

The last two decades years have seen a substantial rise in the field of optogenetics with the discovery and development of light-responsive protein domains that can be genetically incorporated into host cells.^43^ By coupling proteins-of-interest to these light-responsive protein domains, native cellular processes can be studied with greater spatial precision or reengineered to generate novel cellular functions.^44–50^ Numerous light-responsive protein domains have since been discovered with excitation wavelengths ranging from the ultraviolet (∼300 nm) to the near-infrared (>750 nm).^51–57^ To enable spatially resolved activation of split-small GTPases, we chose the improved light-inducible dimerization, iLID, system given the orthogonality of its protein domains to mammalian cells and native chromophore, flavin mononucleotide (FMN).^58^ The availability of iLID variants with different binding affinities in the lit and dark states could allow for tuning of split-small GTPase activation.^58,59^ The iLID system utilizes the light, oxygen, or voltage 2 (LOV2) domain of phototropin 1 from Avena sativa.^60^ Upon exposure to blue light (450 nm – 500 nm), the LOV2 domain undergoes conformational changes by forming a temporary covalent bond between a reactive cysteine residue within the core of the LOV2 protein and FMN. These conformational changes result in the eventual dissociation and unwinding of a C-terminal α-helix, commonly referred to as the Jα-helix. In iLID, the Jα-helix has a short SsrA peptide tag incorporated into its C-terminal end that conditionally binds to its binding partner, SspB, upon blue-light exposure and subsequent uncaging from the core of the LOV2 protein domain. We hypothesized that fusion of our N-terminal and C-terminal split-small GTPase fragments to SspB and iLID, respectively, would allow for light-controlled reassembly of split-small GTPases.

To investigate our hypothesis, we designed light-gated split-Cdc42 (spLIT-Cdc42) constructs by fusing the N-terminal fragment of a constitutively active mutant of Cdc42 (Q61L) to the C-terminus of SspB and the C-terminal fragment to the N-terminus of iLID (**Figure S1**). To monitor expression of our proteins in cells, we fused a red-fluorescent protein to the N-terminus of SspB and a yellow fluorescent protein to the C-terminus of iLID followed by a CAAX box for localization to the cytosolic face of the membrane. Both fragments were subsequently cloned into a vector containing an internal ribosome entry site to allow for bicistronic expression (**Figure S1**; **Tables S1** and **S2**). To explore the tunability of Cdc42 activation, we fused the N-terminal fragment of Cdc42 to the Nano, Milli, and Micro mutants of SspB,^58,59^ yielding three distinct constructs with different binding affinities in the lit and dark state (**Figure S1**). This allowed us to systematically evaluate how the binding affinities of the light-induced dimerization domains influenced downstream Cdc42 signaling outputs.

A well-established, canonical role of Cdc42 signaling in mammalian cells is the induction of filopodia formation.^61^ Filopodia are long, filamentous protrusions from the cell membrane that serve diverse functions in cell movement and intercellular communication.^62^ To first verify the activity of spLIT-Cdc42 in cells, we sought to reproduce previous findings utilizing a rapamycin-gated split-Cdc42 design by globally illuminating cells with blue light (**Figure 2a**).^40^ As a negative control, nontransfected HeLa cells exposed to global illumination with blue light for 30 minutes showed no significant change in filopodia density before or after exposure to light (**Figures 2b** and **2f**). To test our spLIT-Cdc42 constructs, we began with the highest affinity mutant, spLIT-Cdc42-Nano, to increase the likelihood of detecting changes in filopodia density. Transiently transfected HeLa cells were exposed to blue light, imaged (**Figures 2c** and **S2a**), and filopodia formation was quantified demonstrating a 2.9-fold increase in filopodia density after exposure compared to pre-exposure levels (**Figure 2f**). Importantly, only cells expressing both the N- and C-terminal fragments, as determined by imaging the fluorescent protein tags (**Figure S2**), were used in quantification. As an additional control, we mutated the reactive cysteine residue within the LOV2 domain of iLID to an alanine, abolishing the formation of the covalent bond between iLID and FMN upon stimulation with blue light. As expected, illumination of HeLa cells expressing spLIT-Cdc42-Nano C252A did not lead to an increase in filopodia density of the cells (**Figures 2c**, **2f**, and **S2b**). To explore the tunability of spLIT-Cdc42, we then tested the two additional variants of SspB, Micro and Milli. Cells expressing spLIT-Cdc42-Micro exhibited a 2.1-fold increase in filopodia density after illumination (**Figures 2d, 2f**, and **S2c**), albeit with a lower absolute filopodia density compared to cells expressing spLIT-Cdc42-Nano. No change in filopodia density was observed for SspB-Milli, indicating that the lit-state affinity of this iLID system was too weak to drive signaling from the Cdc42 fragments (**Figures 2e, 2f**, and **S2e**). Mutation of the reactive cysteine in spLIT-Cdc42-Micro C252A and spLIT-Cdc42-Milli C252A eliminated any change in filopodia density in transfected cells (**Figures 2d-f**). Taken together, these results demonstrate the ability to tune split-small GTPase signaling responses by modulating the affinity of the light-induced dimerization domains. These results also clearly indicate that spLIT-Cdc42 can optically control Cdc42 activity in mammalian cells enabling the potential investigation of localized signaling effects on cellular phenotypes.

**Figure 2.**
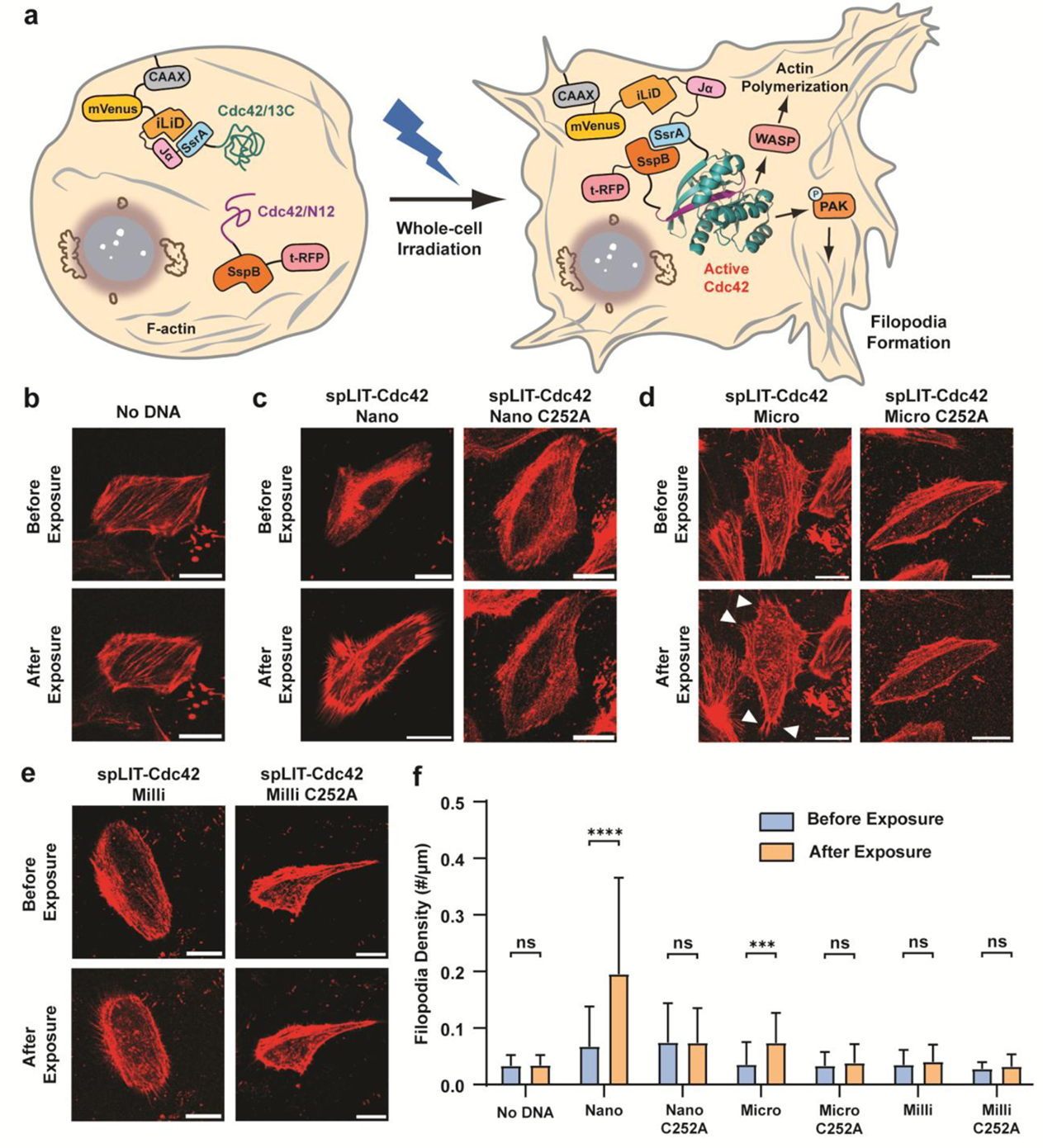
Regulating the formation of filopodia in living cells using spLIT-Cdc42. **a)** Schematic illustrating the whole-cell irradiation of living cells to reassemble spLIT-Cdc42 and induce cell-wide filopodia formation. **b-e)** Representative confocal images (F-actin staining channel shown) before and after whole-cell irradiation of **b)** nontransfected HeLa cells, **c)** HeLa cells expressing the spLIT-Cdc42-Nano systems, **d)** HeLa cells expressing the spLIT-Cdc42-Micro systems, and **e)** HeLa cells expressing the spLIT-Cdc42-Milli systems. Verification of spLIT-Cdc42 fragment expression can be found in **Figure S2** for each cell shown. **f)** Quantification of filopodia density (number of filopodia divided by cell edge length) before and after global irradiation using FiloQuant.^81^ Data for each construct represents the average of at least 30 cells from at least 2 biological replicates. Statistical significance was determined using a two-tailed, unpaired Student’s *t*-test. Error bars represent the standard deviation of the mean. **** indicates a p-value of <0.0001, *** indicates a p-value of <0.001, and ns indicates a p-value of >0.05. All scale bars represent 20 μm. Cells were simultaneously stimulated using 480 nm and 488 nm lasers at 20% intensity (50-55 μW) for 5 minutes followed by a 5-minute rest period in the dark. This was repeated for a total of 30 minutes and cells were immediately imaged.

### Spatial Regulation of Filopodia Formation

We next sought to demonstrate the spatial precision of spLIT-Cdc42 by illuminating only a small subsection of the cell membrane. Using this approach, only protein fragments within the area of exposure should reassemble, leading to a localized cellular response (**Figure 3a**). We began by illuminating one end of a cell expressing spLIT-Cdc42-Nano as verified before light exposure by imaging fluorescent protein tags (**Figures 3b-c** and **S3a**). The change in filopodia density before and after exposure within the illuminated region of the cell (blue square) was then normalized to the change in filopodia density of a nonilluminated region on the opposite end of the cell (gray square). Upon comparison, the illuminated region exhibited a 2.6-fold increase in filopodia density relative to the nonilluminated region (**Figure 3h**). Repeating the experiment with cells expressing the spLIT-Cdc42-Nano C252A mutant (**Figures 3b, 3d**, and **S3b**) resulted in no change in filopodia formation (**Figure 3h**). A similar outcome was observed in cells expressing spLIT-Cdc42-Micro (**Figures 3e-f** and **S3c**) and spLIT-Cdc42-Micro C252A (**Figures 3e, 3g**, and **S3d**) with a relative fold change in filopodia density of 1.4 for spLIT-Cdc42-Micro (**Figure 3h**). These results highlight the ability to spatially control Cdc42 activation using this system and indicate the ability to tune Cdc42 signaling by utilizing iLID systems with varying affinities. Although cells expressing the spLIT-Cdc42-Nano system exhibited higher formation of filopodia when exposed to blue light than those expressing spLIT-Cdc42-Micro, cells expressing the spLIT-Cdc42-Nano system also had a higher level of filopodia formation prior to exposure (**Figures 2f**, **3c**, and **3f**). Therefore, we chose to proceed with the intermediate affinity SspB-Micro iLID system for subsequent experiments, referred to hereafter as the spLIT system, since its reduced dark-state association minimized background activation while still providing a robust light-dependent phenotype. These results demonstrate the ability to spatially control Cdc42 signaling in living cells using the spLIT small GTPase platform.

**Figure 3.**
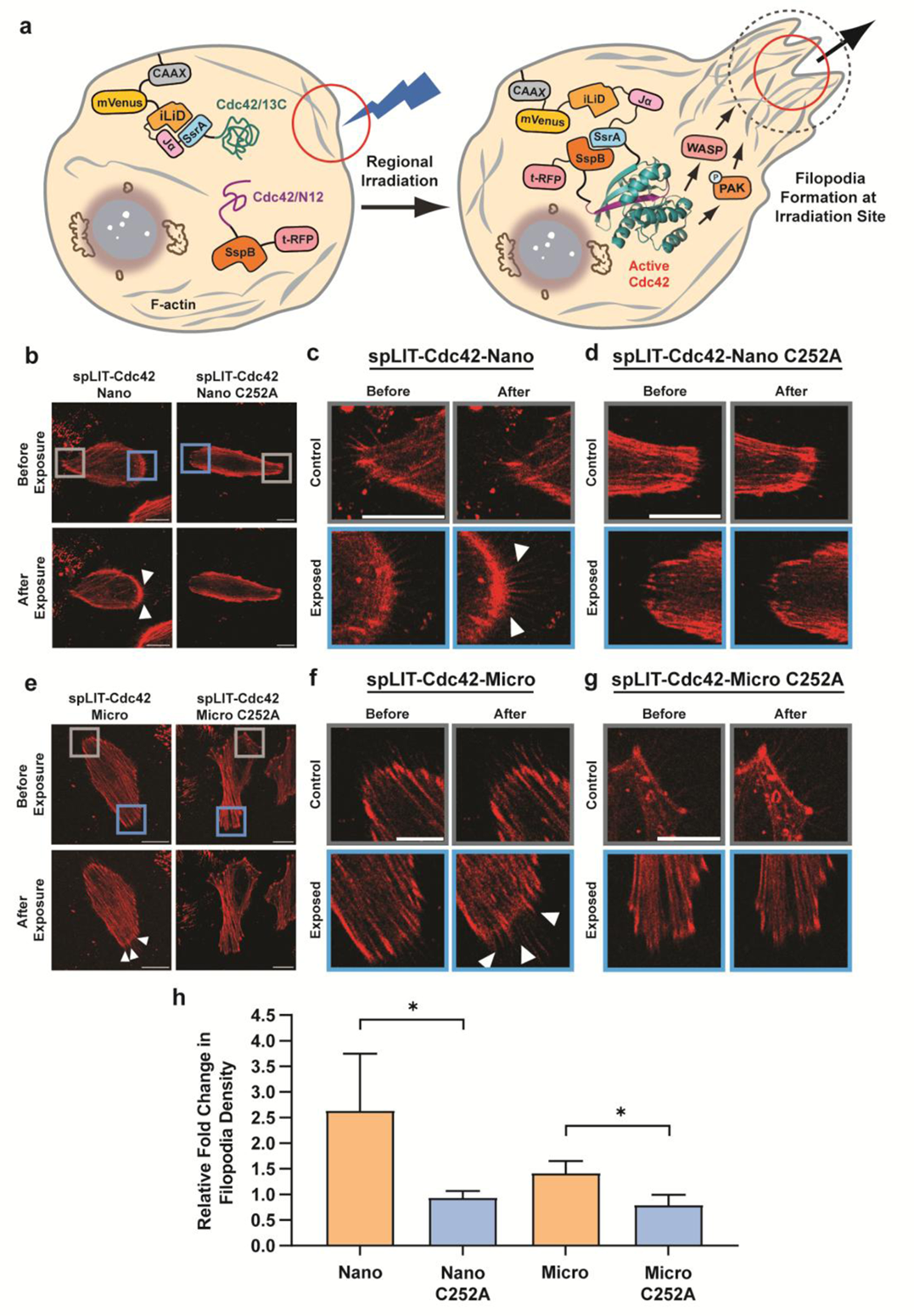
Spatially controlled filopodia formation. **a)** Schematic illustrating the spatially confined activation of filopodia formation via the subcellular activation of spLIT-Cdc42 in living cells. **b)** Representative confocal images (F-actin staining channel shown) of HeLa cells expressing the spLIT-Cdc42-Nano systems before and after site-specific irradiation. Verification of spLIT-Cdc42 fragment expression can be found in **Figure S3** for each cell shown. **c)** Magnified images of the exposed region (blue square) and control region (grey square) of the cell expressing spLIT-Cdc42-Nano from panel b. **d)** Magnified images of the exposed region (blue square) and control region (grey square) of the cell expressing spLIT-Cdc42-Nano C252A from panel b. **e)** Representative confocal images (F-actin staining channel shown) of HeLa cells expressing the spLIT-Cdc42-Micro systems before and after site-specific irradiation. Verification of spLIT-Cdc42 fragment expression can be found in **Figure S3** for each cell shown. **f)** Magnified images of the exposed region (blue square) and control region (grey square) of the cell expressing spLIT-Cdc42-Micro from panel e. **g)** Magnified images of the exposed region (blue square) and control region (grey square) of the cell expressing spLIT-Cdc42-Micro C252A from panel e. **h)** Quantification of the fold-change in filopodia formation before and after irradiation of the exposed end of the cells relative to the nonexposed end of the cells. Data for each construct represents the average of at least 3 cells from independent biological replicates. Statistical significance was determined using a two-tailed, unpaired Student’s *t*-test. Error bars represent standard deviation of the mean. All scale bars represent 20 μm. * indicates a p-value of <0.05. Cells were stimulated using a 480 nm laser at 1-3% intensity (10-15 µW) for 30 seconds followed by a 30-second rest period in the dark. This was repeated for a total of 10 minutes after which cells were imaged.

### Optogenetic Control of Cell Movement Reveals Long-Range Signaling Dynamics

To investigate the modular nature of our optogenetic spLIT-Cdc42 system, we set out to develop spLIT-small GTPase constructs for additional small GTPases. Rac1 regulates the formation of lamellipodia through its effectors, p-21 activating kinase (PAK) and WASP-family verprolin homologous protein (WAVE).^63^ Physiologically, Rac1 aids in the outward projection of the cell membrane at the leading edge, allowing the cell to move forward. We hypothesized that replacing the Cdc42 fragments within spLIT-Cdc42 with the corresponding Rac1 fragments would lead to the localized formation of cell membrane protrusions and the directional control of cell movement (**Figure 4a**). Accordingly, we replaced the Cdc42 fragments of spLIT-Cdc42 and spLIT-Cdc42 C252A with the corresponding fragments of Rac1, generating spLIT-Rac1 and spLIT-Rac1 C252A, respectively (**Figure S1**; **Tables S1** and **S2**). Excitingly, spatially restricted exposure of MEF cells expressing spLIT-Rac1 to light led to clear protrusion of the cell membrane in the direction of light exposure (**Figures 4b** and **S4a**; **Videos S1-S4**). Conversely, cells expressing the spLIT-Rac1 C252A mutant did not respond to light stimulation (**Figures 4c** and **S5b**; **Videos S5-S9**). Since cells expressing the spLIT-Rac1 construct did not move at a consistent rate from cell-to-cell, we sought to quantify movement across multiple cells using an index-based approach. In short, individuals with varying levels of familiarity with the experiment rated the directional movement of each cell relative to an arbitrary red dot placed on a video of each cell (**Figure 4d**). Importantly, raters were unaware of which protein construct each cell expressed nor the position of light exposure. Aggregated ratings revealed a clear trend: cells expressing spLIT-Rac1 predominately moved towards the direction of light exposure while cells expressing spLIT-Rac1 C252A exhibited a slight retraction away from the light stimulus (**Figure 4e**). These data indicate the ability to control the direction of cell movement using spLIT-Rac1.

**Figure 4.**
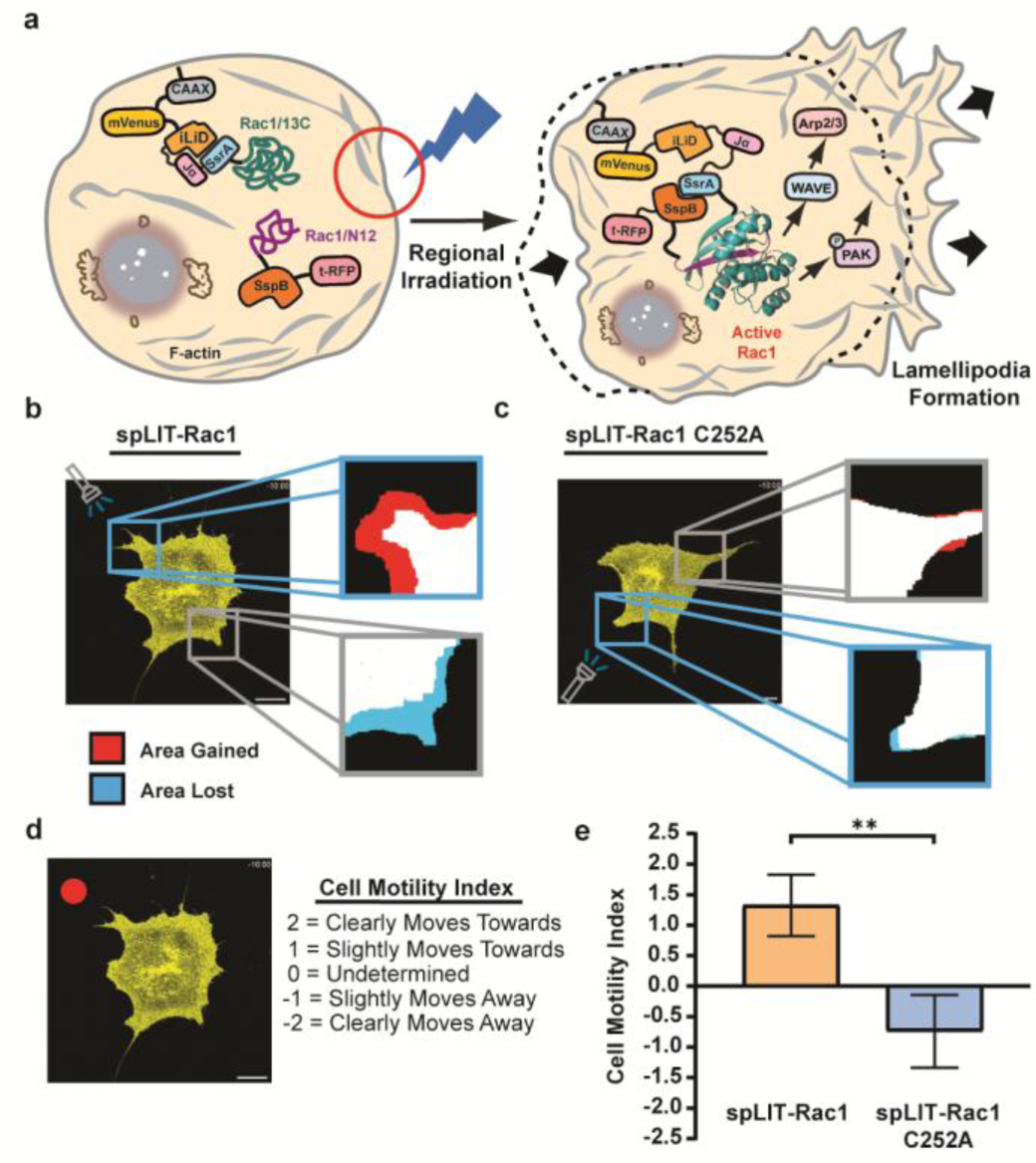
Controlling directional cell movement via spatially restricted Rac1 activation. **a)** Schematic illustrating the spatial regulation of lamellipodia formation via the subcellular activation of spLIT-Rac1 in living cells. **b)** Representative confocal image (mVenus channel shown) of a MEF cell expressing spLIT-Rac1 and selectively exposed to light. Verification of spLIT-Rac1 fragment expression can be found in **Figure S4a**. Zoomed in images illustrate the change in cell area post exposure within the exposed region (blue square) and control region (grey square). **c)** Representative confocal image (mVenus channel shown) of a MEF cell expressing spLIT-Rac1 C252A and selectively exposed to light. Verification of spLIT-Rac1 C252A fragment expression can be found in **Figure S4b**. Zoomed in images illustrate the change in cell area post exposure within the exposed region (blue square) and control region (grey square). **d)** Left - Representative image of a video given to raters. The red dot is a reference point. Right - Raters were tasked with choosing one of the “Cell Motility Index” rankings for each cell. **e)** Quantification of cell motility for MEF cells expressing spLIT-Rac1 (n=4) and spLIT-Rac1 C252A (n=5). Each cell is from an independent biological replicate. Data were aggregated from 13 independent, blinded ratings for each cell. Statistical significance was determined using a two-tailed, unpaired Student’s *t*-test. Error bars represent standard deviation of the mean. All scale bars represent 20 μm. ** indicates a p-value of <0.01. Cells were stimulated using a 480 nm laser at 1-3% intensity (10-15 µW) for 30 seconds, followed by a 30-second rest period in the dark. This was repeated for a total of 10 minutes after which cells were imaged.

Interestingly, we also observed membrane retraction on the non-exposed edge of cells expressing spLIT-Rac1 which was consistent in magnitude to the protrusions on the exposed edge, leading to observable cell motion in the direction of light exposure (**Figures 4b** and **S6**; **Videos S1-S4**). This phenomenon was not observed in cells expressing spLIT-Rac1 C252A (**Figure 4c**; **Videos S5-S9**). Previous work utilizing a photoactivatable, full-length Rac1 protein (PA-Rac1) demonstrated similar results upon spatially restricted exposure of cells with blue light and is attributed to the dynamic, long-range regulation of RhoA activity by Rac1.^35^ These results highlight the modularity of the spLIT small GTPase platform, demonstrating that split-small GTPase fragments can be interchanged without the need for case-by-case optimization. Moreover, the spLIT-Rac1 system validates the ability to identify long-range signaling effects resulting from direct, local activation of a small GTPase.

### Uncovering Long-Range RhoA Signaling

While the long-range effects from local activation of Rac1 signaling have previously been established,^35^ long-range effects of localized RhoA signaling (i.e. the potential propagation of RhoA activity across the cell) are less well studied due in part to the lack of optogenetic tools to directly activate RhoA with subcellular precision. RhoA regulates downstream effectors such as Rho-associated kinase (ROCK) and mDia1, leading to actomyosin contractility, stress fiber formation, and focal adhesion assembly.^64^ These processes contribute to large-scale contraction of the cell membrane, allowing for force generation and directional cell movement. Previous work using optogenetic systems capable of localizing a RhoAGEF to different subcellular regions demonstrated the ability to spatially control RhoA-induced cell membrane retraction.^65^ However, a subsequent study utilizing a similar optogenetic RhoAGEF system highlighted the inherent cross-talk associated with GEF-mediated small GTPase activation, yielding the opposite phenotype (protrusion versus retraction) resulting from simultaneous activation of Cdc42.^66^ Hence, a direct approach to locally active RhoA is need to unambiguously attribute signaling changes to this small GTPase. Intriguingly, implementation of an optogenetic method to release constitutively active RhoA or ROCK from the mitochondria demonstrated long-range effects of RhoA signaling on axon formation in neurons, possibly through the diffusion of RhoA effectors, such as ROCK,^67^ and/or through the inherent feed-forward control of RhoA activity.^68^ Additionally, it has been reported that knockdown of RhoA activity in human cancer cells leads to simultaneous lateral expansion of the cell membrane at opposing ends of the cell.^69^ However, global knockdown of protein expression can potentially obscure effects from spatially localized signaling. Given taht direct evdience for the role of RhoA in long-range signaling has remaing elusive, we sought to investigate whether direct, spatially localized activation of spLIT-RhoA could induce long-range signaling effects on cell membrane retraction in living cells (**Figure 5a**).

**Figure 5.**
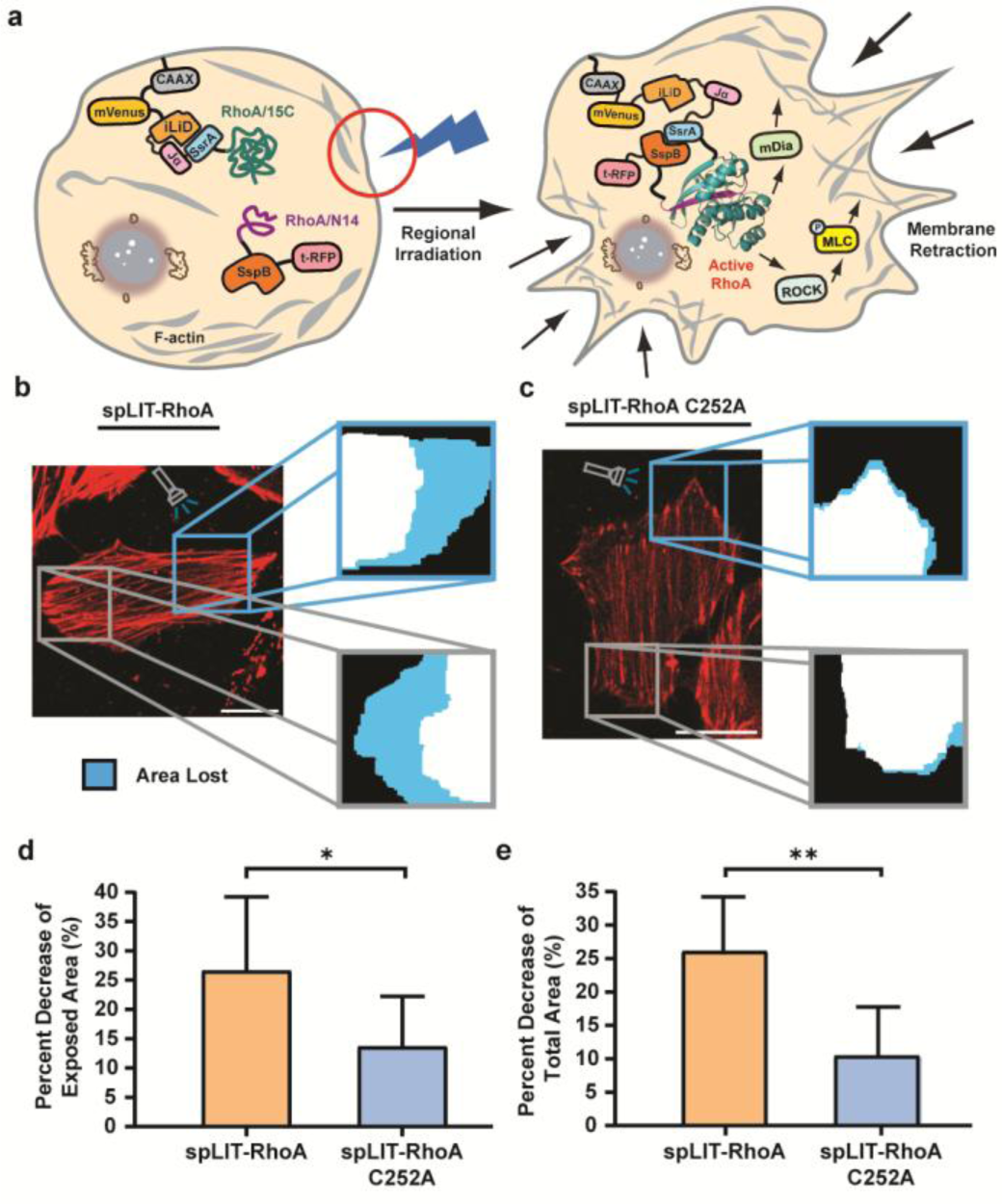
Localized activation of spLIT-RhoA leads to bipolar cell retraction. **a)** Schematic illustrating the spatially confined activation of spLIT-RhoA in living cells. **b)** Representative confocal image (F-actin staining channel shown) of a HeLa cell expressing spLIT-RhoA and exposed to light on one side (blue square). Verification of spLIT-RhoA fragment expression can be found in **Figure S5a** for the cell shown. Zoomed in images illustrate the reduction in cell area post exposure within the exposed region (blue square) and control region (grey square). **c)** Representative confocal image (F-actin staining channel shown) of a HeLa cell expressing spLIT-RhoA C252A and exposed to light on one side (blue square). Verification of spLIT-RhoA C252A fragment expression can be found in **Figure S5b** for the cell shown. Zoomed in images illustrate the cell area reduction post exposure within the exposed region (blue square) and control region (grey square). **d)** Quantification of the reduction in cell area within the exposed region for cells expressing spLIT-RhoA and spLIT-RhoA C252A. Data for each construct represents the average of at least 6 cells from at least 5 independent biological replicates. **e)** Quantification of the decrease in total area of the same cells expressing spLIT-RhoA (n=6) or spLIT-RhoA C252A (n=9) post irradiation from panel d. HeLa cells expressing the active spLIT-RhoA system exhibit a larger decrease in total cell area post irradiation. Statistical significance was determined using a two-tailed, unpaired Student’s *t*-test. Error bars represent standard deviation of the mean. All scale bars represent 20 μm. * indicates a p-value of <0.05. Cells were stimulated using a 480 nm laser at 1-3% intensity (10-15 µW) for 30 seconds followed by a 30-second rest period in the dark. This was repeated for a total of 10 minutes.

We previously demonstrated that CIDs could be used to gate split-RhoA-mediated cell membrane retraction on a global scale within cells.^40,41^ To test whether spatially precise activation of RhoA signaling could induce long-range signaling in living cells, we cloned spLIT-RhoA and spLIT-RhoA C252A (**Figure S1**; **Tables S1** and **S2**). Spatially restricted illumination of HeLa cells expressing spLIT-RhoA (**Figures 5b** and **S5a**; **Video S10**) led to a 2-fold increase in retraction compared to cells expressing spLIT-RhoA C252A (**Figures 5c-d** and **S5b**; **Video S11**), indicating the ability to activate RhoA signaling with light. Interestingly, activation of spLIT-RhoA led to an equivalent decrease in cell area at both the exposed and non-exposed poles of cells (**Figure 5b**). However, a large decrease in cell area in either the exposed or control poles of cells expressing spLIT-RhoA C252A was not observed (**Figure 5c**). Since the retraction of the cell membrane in the non-exposed region was equivalent in magnitude to the exposed region for spLIT-RhoA expressing cells, the difference in the observed decrease in cell area of the exposed regions (**Figure 5d**) was comparable to the difference in global decrease in cell area between spLIT-RhoA expressing cells and spLIT-RhoA C252A expressing cells (2.6-fold higher for cells expressing spLIT-RhoA [**Figure 5e**]). Importantly, a pronounced decrease in cell area was not observed in either exposed or control poles of spLIT-RhoA C252A expressing cells (**Figures 5c-e**), confirming that bipolar retraction arises specifically from RhoA activation rather than optical stimulation. These results provide direct evidence that local RhoA activation can elicit cooridnated, long-range contractile reponsese across the cell, culminating in a striking bipolar membrane retraction phenotype. This finding establishes spLIT-RhoA as a powerful tool for dissecting mechanical signal propogation within cells and suggests that localized RhoA activity can raplidly coordinate cytoskeletal tension across distant cellular domains. Ongoing efforts in our lab are focused on defining the mechanism of long-range RhoA signaling in cells.

## Conclusion

In summary, we introduce the first modular optogenetic platform that enables direct light-controlled activation of small GTPases in living cells with high spatial resolution. We engineered spLIT-small GTPase constructs for constitutively active forms of Cdc42, Rac1, and RhoA without the need for case-by-case optimization, achieving localized reconstitution and functional activation within living cells. This method allowed us to directly manipulate complex cellular behaviors, including filopodia formation and directed cell migration. Together, these results establish spLIT-small GTPases as a powerful plug-and-play platform for dissecting spatial signaling dynamics in live cells.

Notably, localized activation of spLIT-Rac1 not only leads to the formation of cell membrane protrusions at the site of light exposure but also leads to retraction of the cell membrane at the opposing edge of the cell (**Figures 4b** and **S6**; **Videos S1-4**). In separate experiments, spatial activation of spLIT-RhoA enabled observation of bipolar membrane retraction (**Figure 5b**; **Video S10**), a previously undisclosed long-range effect of direct, localized RhoA activation. It is well established that migrating cells typically maintain transient, isolated clusters of active RhoA at their leading edge and a concentrated pool of active RhoA at their trailing edge to aid in contraction of actomyosin and retraction of the cell membrane during cell motion,^70–74^ yet direct evidence for long-range RhoA signaling has been lacking. Previous work has highlighted the context-dependent mutual regulation of Rac1 and RhoA, indicating the ability of Rac1 to inhibit and/or activate RhoA.^75–79^ Our data provide the first demonstration that localized RhoA activation alone can propagate contractile signaling to distant regions of the same cell, highlighting a potential mechanism by which Rac1-triggered local RhoA activation could facilitate rear-edge retraction during directed migration. Current efforts in our lab are focused on further dissecting the molecular basis of this long-range RhoA signaling propagation.

Although binding of the iLID system is rapidly reversible (reversion t_1/2_ = 18s),^58,59,80^ sustained signaling from spLIT-small GTPases can be observed up to 30-45 minutes after illumination (**Videos S1-4** and **S10**). This is likely due to the extensive contacts formed between the N12 and 13C small GTPase fragments upon reassembly (**Figure S7**). Our lab is currently investigating mutations within the N12 fragment that may yield more readily reversible systems.

Looking ahead, the spLIT-small GTPase system provides a potentially powerful tool to explore the spatial regulation of small GTPase signaling (e.g. long-range RhoA signaling induced by localized RhoA activation). Pairing split-small GTPase fragments with spectrally orthogonal light-induced dimerization domains^51–57^ may enable multiplexed activation of different small GTPases. Thus, we believe that the technology described here will find broad applications in studying fundamental aspects of small GTPase signaling and in synthetic biology efforts to engineer new functions into living systems.

## Methods

### DNA Vector Synthesis

Individual DNA segments were amplified using conventional PCR DNA amplification and fused together using overlap extension PCR. The split-small GTPase sequences and mVenus sequence were obtained from previously published plasmids.^40,41^ The tagRFPt-SspB Nano sequence was obtained from pLL7.0: tgRFPt-SSPB WT (Addgene: 60415). The tagRFPt-SspB Micro sequence was obtained from pLL7.0: tgRFPt-SSPB R73Q (Addgene: 60416). The iLID sequence was obtained from pLL7.0: Venus-iLID-CAAX (from KRas4B) (Addgene: 60411). The SspB Milli sequence was obtained from pAAV hSyn mCh-IRES-SspB(milli)-BoNT/B(147-441, Y365A) (Addgene: 122985). Final DNA constructs were ligated into an empty pIRES vector using restriction enzyme-based techniques. Prep kits from Qiagen were used for plasmid preparation. Site-directed mutagenesis was performed using the QuikChange II XL Site-Directed Mutagenesis Kit (Agilent Technologies). Sequence integrity was verified using Sanger Sequencing (Genewiz).

### Mammalian Cell Culture

HeLa cells (ATCC, CCL-2) and MEF cells (ATCC, C57BL/6) were used in this work. Stocks were kept frozen in liquid nitrogen until use. Mammalian cells were cultured at 37°C and 5% CO_2_ for 72-96 hours before passaging. Cell culture medium, DMEM (Dulbecco’s Modified Eagle Medium, Thermo Fisher), was supplemented with 10% (v/v) fetal bovine serum (Thermo Fisher),100 U/ml penicillin and 100 g/ml streptomycin (Thermo Fisher). To passage, cells were washed in 5 mL of DPBS (Thermo Fisher), suspended using 2 mL of TrypLE™ Express Enzyme (Thermo Fisher) for 10 minutes, centrifuged at 100g for 5 minutes at 4°C, resuspended and diluted (1:10) in prewarmed DMEM.

### Mammalian Cell Transfection

Cells were plated at an initial confluency of 20% using a Countess™ 3 automated cell counter (Invitrogen) and cultured in DMEM at 37°C with 5% CO_2_ for 24 hours. Cells were then transfected with DNA vectors and incubated overnight. Cells were transfected using Lipofectamine™ 3000 (Thermo Fisher) based on the manufacturer’s protocol. Once transfected, HeLa and MEF cells were covered in aluminum foil and cultured overnight at 37°C with 5% CO_2_ before imaging.

### Imaging Mammalian Cell Lines

Prior to imaging, HeLa cells were stained with 1 µL per dish of CellMask™ Deep Red Actin Tracking Stain (Thermo Fisher) to monitor F-actin dynamics. Importantly, staining with F-actin dye post-transfection was performed in a dark room under red light to avoid incidental exposure to blue light. Additionally, microscope computer screens were covered with yellow long-pass filters to avoid exposure to blue light. Expression of tagRFP-T was monitored by excitation at 555 nm and emission from 570 nm - 610 nm. Expression of mVenus was monitored by excitation at 514 nm and emission from 520 nm - 550 nm. Cells stained with the CellMask™ Deep Red Actin Tracking Stain were imaged using an excitation wavelength of 652 nm and emission detection wavelengths from 670 nm - 710 nm. Confocal images were obtained on a Leica STELLARIS 8 confocal/FLIM/tauSTED microscope system equipped with a tunable white light laser. Cells were imaged using the HC PL APO 40x/1.30 OIL CS2 objective lens (Leica Microsystems), and the confocal microscope was controlled using the LAS X software. All images were processed using the open-source software ImageJ (Fiji). The scale bar in all images is 20 µm.

### Global Activation of spLIT-Cdc42

To activate spLIT-Cdc42, laser settings and cell exposure times to blue light were adapted from prior literature^58^ and optimized to improve signal readout while limiting cell toxicity. Dual wavelengths of 488 nm and 480 nm were used to activate the iLID system and drive reassembly of the Cdc42 fragments. The laser lines for both wavelengths used to activate the iLID system were set to 85% power and 20% intensity (25-27 µW each). The ROI was set to the maximum to distribute the laser light across the entire dish. Prolonged exposure to blue light can lead to DNA damage and induction of apoptosis in cells. As such, utilizing blue light to control protein activity may cause off-target effects that could impact results in certain applications, such as the study of apoptotic signaling pathways. To combat this issue, we limited exposure to blue-light to short, non-continuous bursts. Cells were exposed to the blue laser light for 5 minutes followed by 5 minutes in darkness to activate the iLID system. The cells were then imaged using the F-actin stain and fluorescent protein channels to monitor any changes in filopodial formation. This process was repeated 3 times for a total time of 30 minutes.

### Local Activation of spLIT-small GTPases

For local activation of spLIT-Cdc42, an ROI was drawn on a subsection of a single, transfected cell using the LAS X software. The laser settings and cell exposure times were optimized to improve signal readout while limiting cell toxicity. A 480 nm laser (set to 85% power and 1-3% intensity [10-15 µW]) was used to activate the iLID system within the ROI. The ROI was illuminated for 30 seconds, followed by 30 seconds of darkness. This process was repeated for a total of 10 minutes. Images were collected immediately before illumination and immediately after illumination. During image processing, a control ROI of the exact same size was drawn on the opposite side of the cell that was illuminated and was used as an internal control.

For local activation of spLIT-RhoA and spLIT-Rac1, an ROI was drawn on a subsection of a single, transfected cell using the LAS X software. Again, the laser settings and cell exposure times were optimized to improve signal readout while limiting cell toxicity. A 480 nm laser (set to 85% power and 1-3% intensity [10-15 µW]) was used to activate the iLID system within the ROI. The ROI was illuminated for 30 seconds, followed by 30 seconds of darkness. This process was repeated for a total of 10 minutes. Images were collected immediately before illumination, immediately after illumination, and every 5 minutes after illumination. During image processing, a control ROI of the exact same size was drawn on the opposite side of the cell that was illuminated and was used as an internal control for split-small GTPase activation.

### Data Processing and Statistical Analysis

To quantify Cdc42-induced filopodia formation, we utilized the open-source ImageJ plugin, FiloQuant.^81^ This software collects data on various cell-surface parameters, including the number and length of filopodia and the cell edge length.^82^ We determined the density of detected filopodia per cell by dividing the number of filopodia of a cell by the cell edge length to gauge the influence of our spLIT-Cdc42 system on filopodia formation. For each plasmid vector tested, the change in filopodia density from at least 30 transfected cells was aggregated and quantified. In each experiment, cells were collected from multiple, independently treated dishes.

To quantify RhoA-induced cell retraction, the area for each cell was determined using ImageJ. For each spLIT-RhoA construct, the cell membrane retraction of at least 7 cells was aggregated and analyzed. In each experiment, cells were collected from multiple, independently treated dishes.

For evaluation of spLIT-Rac1 induced signaling, images were collected pre-light exposure and every 5 minutes post-light exposure. For spLIT-Rac1, 4 cells were analyzed from independently cultured dishes. For spLIT-Rac1 C252A, 5 cells were analyzed from independently cultured dishes. All images for each individual cell were then compiled into a short video using ImageJ. For quantification, 13 individual raters were given the video for each cell and tasked with scoring the movement of each cell relative to a red dot placed on the video. Specifically, each rater selected one of the following prompts for each cell: “-2: Clearly Moves Away”; “-1: Slightly Moves Away”, “0: Undetermined”; “1: Slightly Moves Towards”; “2: Clearly Moves Towards”. Importantly, each rater did not know that the red dot indicated the direction of light exposure and did not know which construct each cell was expressing. All 13 ratings for each cell were averaged and then aggregated according to the construct expressed. Each cell was collected from an independently treated dish.

All statistical analyses were conducted using unpaired, two-tailed Student’s t-tests. All error bars represent the standard deviation of the mean. ns: non-significant, p-value ≥ 0.05; *: p-value < 0.05; **: p-value < 0.01; ***: p-value < 0.001; ****: p-value < 0.0001.

## Supporting Information

Supporting Figures, Tables, Video Captions, and Instrumentation and Methods

## Acknowledgements

We thank the W.M. Keck Center for Cellular Imaging for the usage of the Leica STELLARIS 8 confocal/FLIM/tauSTED microscope system (NIH OD030409) and members of the C. Stains lab for editing the manuscript and helpful conversations. We acknowledge financial support from the NIH (R35GM148221) and the University of Virginia. This research was supported by state funding within the University of Virginia Comprehensive Cancer Center. The content of this work is solely the responsibility of the authors and does not necessarily represent the official views of the NIH.

## TOC Figure

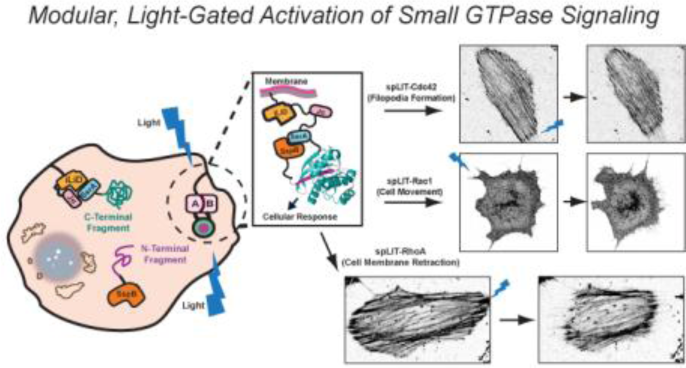

