## Supplementary Material for "A Modular Platform for the Optogenetic Control of Small GTPase Activity in Living Cells Reveals Long-Range RhoA Signaling"

### Figure S1

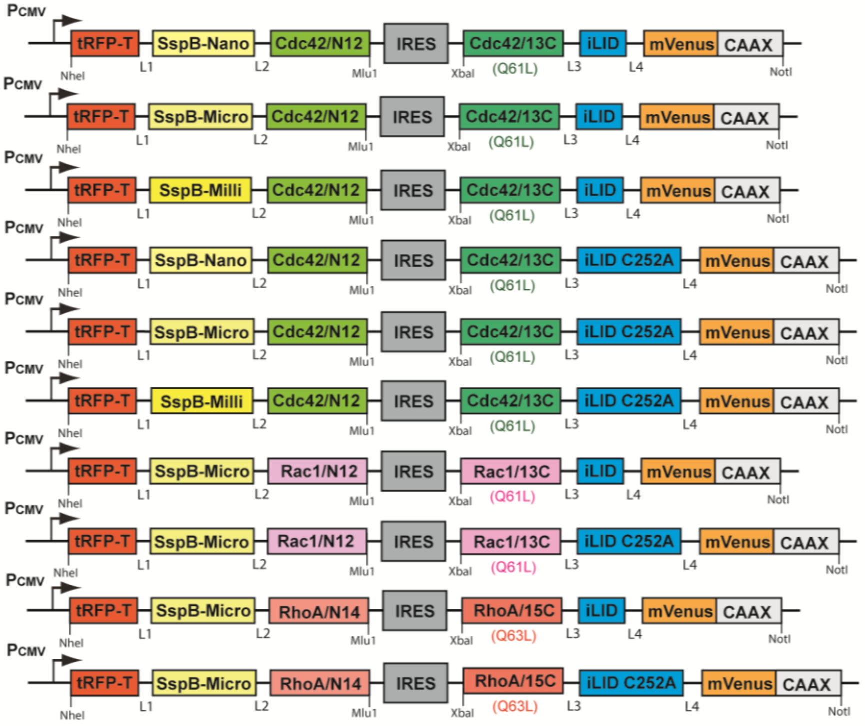

L1: SGLRSRAQASNEFGIDLSGLTLQEF
L2: QISYASRGGGSSGGGEL
L3: GDPNWELVYTARLQGGGSSGGGQISYASRG
L4: GSGSGS
CAAX: KKKKKKSKTKCVIM

A schematic of the protein constructs used for mammalian expression. IRES is an internal ribosomal entry site. Q61L and Q63L denote constitutively active small GTPase mutants. Complete sequence information is provided in **Table S2**.

### Figure S2

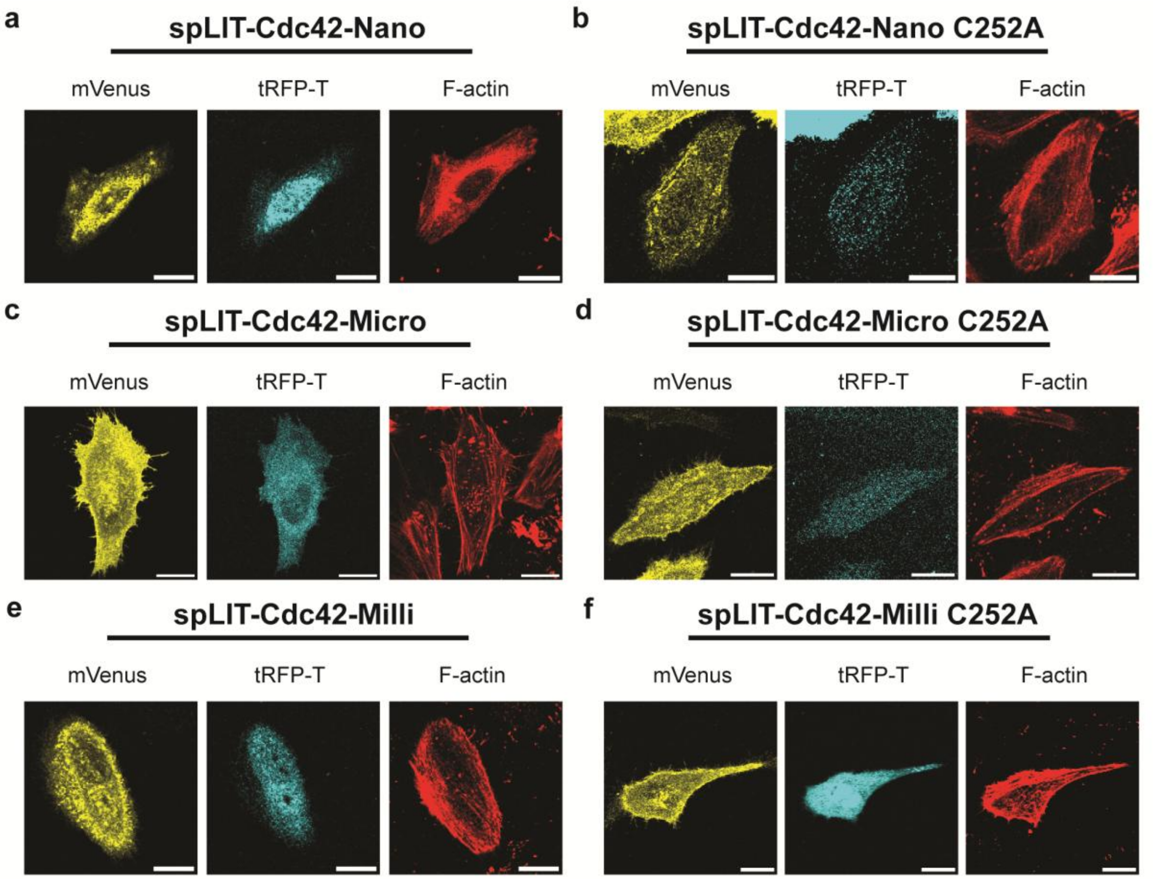

Representative multichannel confocal images of the HeLa cells from **Figure 2** expressing (a) spLIT-Cdc42-Nano, (b) spLIT-Cdc42-Nano C252A, (c) spLIT-Cdc42-Micro, (d) spLIT-Cdc42-Micro C252A, (e) spLIT-Cdc42-Milli, and (f) spLIT-Cdc42-Milli C252A. mVenus fluorescence confirms the expression of the C-terminal fragment and tRFP-T fluorescence confirms the expression of the N-terminal fragment. Scale bar in all images represents 20 µm.

### Figure S3

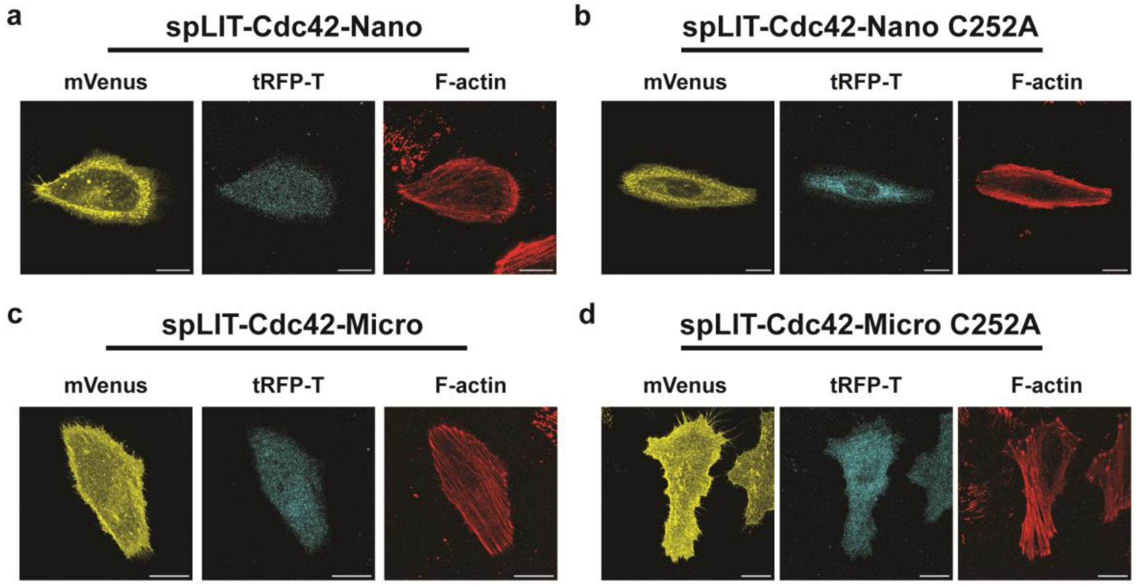

Representative multichannel confocal images for the HeLa cells in **Figure 3** expressing (a) spLIT-Cdc42-Nano, (b) spLIT-Cdc42-Nano C252A, (c) spLIT-Cdc42-Micro, and (d) spLIT-Cdc42-Micro C252A. mVenus fluorescence confirms the expression of the C-terminal fragment and tRFP-T fluorescence confirms the expression of the N-terminal fragment. Scale bar in all images represents 20 μm.

**Figure S4**

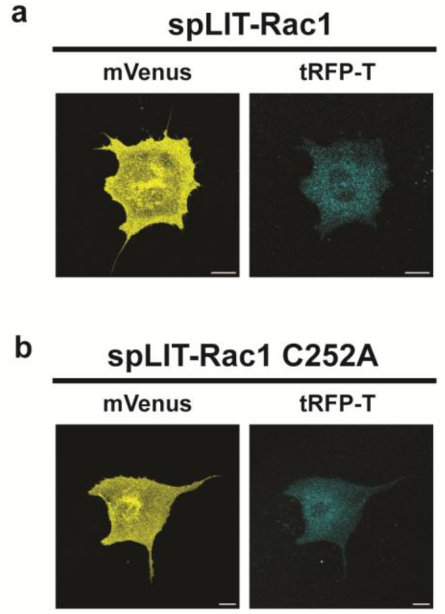

Images of HeLa cells expressing spLIT-Rac1 and spLIT-Rac1 C252A constructs. (a) Representative multichannel confocal images for the MEF cell in **Figure 4b**. (b) Representative multichannel confocal images for the MEF cell in **Figure 4c**. mVenus fluorescence confirms the expression of the C-terminal fragment and tRFP-T fluorescence confirms the expression of the N-terminal fragment. Scale bar in all images represents 20 μm.

### Figure S5

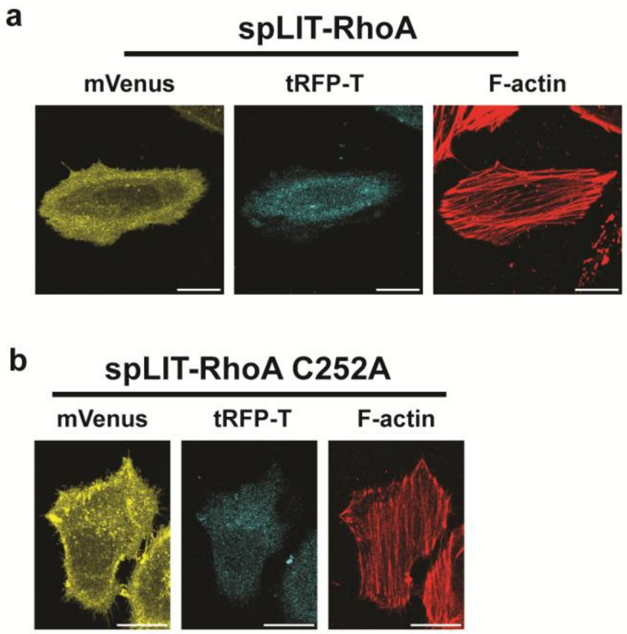

Images of HeLa cells expressing spLIT-RhoA and spLIT-RhoA C252A constructs. (a) Representative multichannel confocal images for the HeLa cell in **Figure 5b**. (b) Representative multichannel confocal images for the HeLa cell in **Figure 5c**. mVenus fluorescence confirms the expression of the C-terminal fragment and tRFP-T fluorescence confirms the expression of the N-terminal fragment. Scale bar in all images represents 20 μm.

**Figure S6**

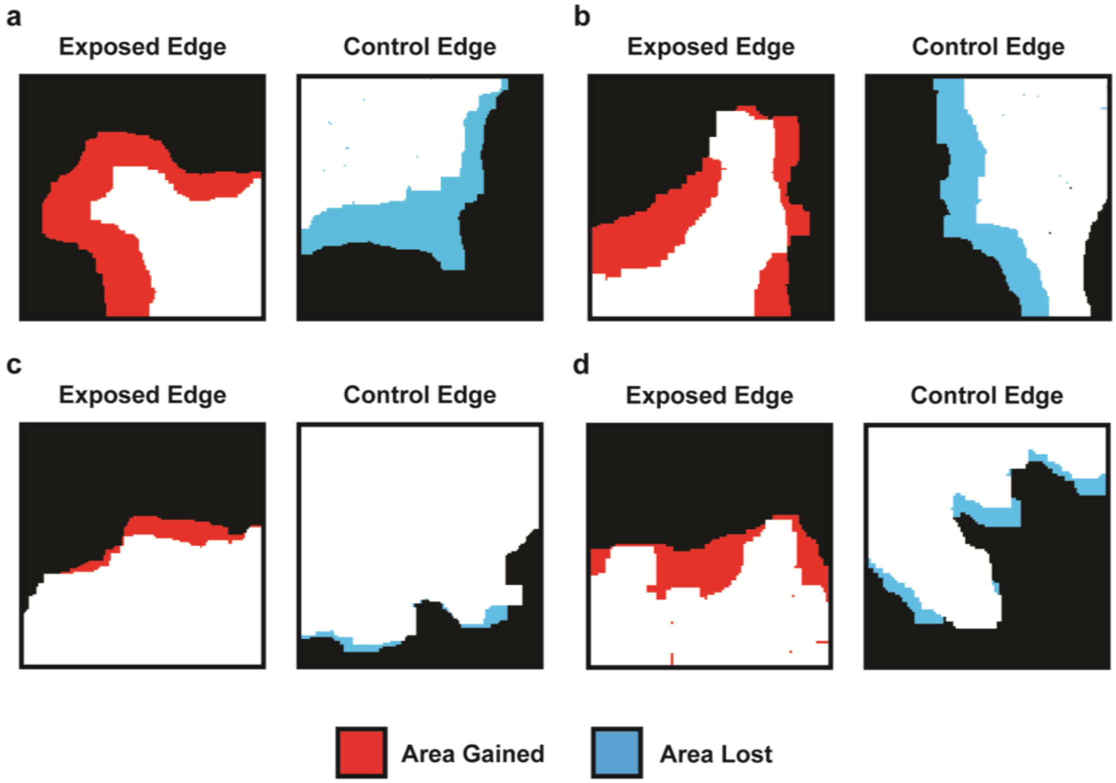

Heat maps illustrate retraction of the control edge of cells expressing spLIT-Rac1 upon irradiation of the exposed edge to blue light. (a) Heat map for the cell in **Video S1**. (b) Heat map for the cell in **Video S2**. (c) Heat map for the cell in **Video S3**. (d) Heat map for the cell in **Video S4**. White regions indicate the starting point for the cells before exposure to blue light. Red regions indicate areas of cell membrane protrusion post-exposure. Blue regions indicate areas of cell membrane retraction post-exposure.

**Figure S7**

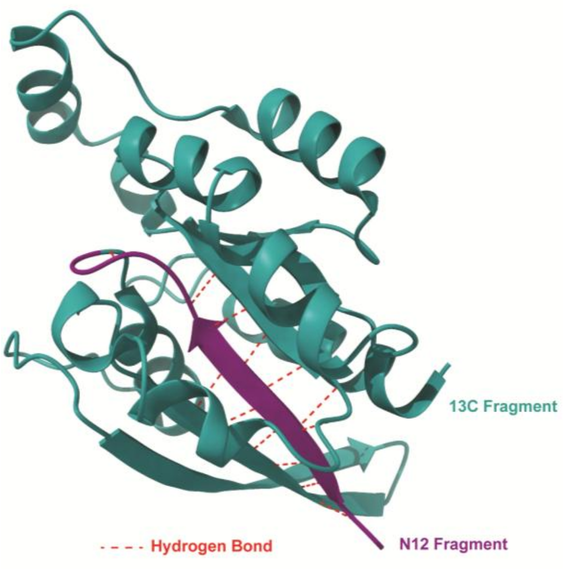

Crystal structure of the small GTPase Cdc42 (PDB: 2ODB). The purple region highlights the N12 fragment of Cdc42. The cyan region highlights the 13C fragment of Cdc42. The red dashed lines indicate hydrogen bonds formed between the N12 and 13C fragments upon reassembly and reconstitution of full-length Cdc42.

### Supplemental Video Captions

**Supplemental Videos 1-4.** Videos of MEF cells expressing active spLIT-Rac1. Upon irradiation, each cell gradually moves in the direction of light exposure (indicated by the blue circle in the first frame). The time for each frame is indicated in the top right corner. The scale bar represents 20 μm.

**Supplemental Videos 5-9.** Videos of MEF cells expressing inactive spLIT-Rac1 C252A. Upon irradiation, each cell either exhibits no directional movement or begins to slightly move away from the direction of light exposure (indicated by the blue circle in the first frame). The time for each frame is indicated in the top right corner. The scale bar represents 20 μm.

**Supplemental Video 10.** A representative video of a HeLa cell expressing active spLIT-RhoA. Upon irradiation, the cell membrane retracts inwards away from the direction of light exposure (indicated by the blue circle in the first frame). The time for each frame is indicated in the top right corner. The scale bar represents 20 μm.

**Supplemental Video 11.** A representative video of a HeLa cell expressing inactive spLIT-RhoA C252A. Upon irradiation, the cell membrane exhibits little movement away from the direction of light exposure (indicated by the blue circle in the first frame). The time for each frame is indicated in the top right corner. The scale bar represents 20 μm.

**Table S1. Construct descriptions and Addgene IDs.**

| Construct | Description | Addgene ID |
|---|---|---|
| spLIT-Cdc42-Nano | pIRES-tagRFPt-SspB Nano-Cdc42/N12-Cdc42/13C-iLID-mVenus-CAAX | 243661 |
| spLIT-Cdc42-Micro | pIRES-tagRFPt-SspB Micro-Cdc42/N12-Cdc42/13C-iLID-mVenus-CAAX | 243662 |
| spLIT-Cdc42-Milli | pIRES-tagRFPt-SspB Milli-Cdc42/N12-Cdc42/13C-iLID-mVenus-CAAX | 243663 |
| spLIT-Cdc42-Nano C252A | pIRES-tagRFPt-SspB Nano-Cdc42/N12-Cdc42/13C-iLID C252A-mVenus-CAAX | 243664 |
| spLIT-Cdc42-Micro C252A | pIRES-tagRFPt-SspB Micro-Cdc42/N12-Cdc42/13C-iLID C252A-mVenus-CAAX | 243665 |
| spLIT-Cdc42-Milli C252A | pIRES-tagRFPt-SspB Milli-Cdc42/N12-Cdc42/13C-iLID C252A-mVenus-CAAX | 243666 |
| spLIT-Rac1 | pIRES-tagRFPt-SspB Micro-Rac1/N12-Rac1/13C-iLID-mVenus-CAAX | 243667 |
| spLIT-Rac1 C252A | pIRES-tagRFPt-SspB Micro-Rac1/N12-Rac1/13C-iLID C252A-mVenus-CAAX | 243668 |
| spLIT-RhoA | pIRES-tagRFPt-SspB Micro-RhoA/N14-RhoA/15C-iLID-mVenus-CAAX | 243669 |
| spLIT-RhoA C252A | pIRES-tagRFPt-SspB Micro-RhoA/N14-RhoA/15C-iLID C252A-mVenus-CAAX | 243670 |

**Table S2. Sequences for expression constructs.**

**1) pIRES-tagRFPt-SspB Nano-Cdc42/N12-Cdc42/13C-iLID-mVenus-CAAX**

Amino Acid Sequence:

MVSKGEELIKENMHMKLYMEGTVNNHHFKCTSEGEGKPYEGTQTMRIKVVEGGPLPFAFDILA
TSFMYGSRTFINHTQGIPDFFKQSFPEGFTWERVTTYEDGGVLTATQDTSLQDGCLIYNVKIRG
VNFPSNGPVMQKKTLGWEANTEMLYPADGGLEGRTDMALKLVGGGHLICNFKTTYRSKKPAK
NLKMPGVYYVDHRLERIKEADKETYVEQHEVAVARYCDLPSKLGHKLNGMDELYKSGLRSRAQ
ASNEFGIDLSGLTLQEFSSPKRPKLLREYYDWLVDNSFTPYLVVDATYLGVNVPVEYVKDGQIVL
NLSASATGNLQLTNDFIQFNARFKGVSRELYIPMGAALAIYARENGDGVMFEPEEIYDELNIGQIS
YASRGGGSSGGGELQTIKCVVVGDGA* **---IRES Region---**
MGVGKTCLLISYTTNKFPSEYVPTVFDNYAVTMIGGEPYTLGLFDTAGLEDYDRLRPLSYPQT
DVFLVCFSVVSPSSFENVKEKWVPEITHHCPKTPFLLVGTQIDLRDDPSTIEKLAKNKQKPITPET
AEKLARDLKAVKYVECSALTQRGLKNVFDEAILAALEPPETQPGDPNWELVYTARLQGGGSSG
GGQISYASRGEFLATTLERIEKNFVITDPRLPDNPIIFASDSFLQLTEYSREEILGRNCRFLQGPET
DRATVRKIRDAIDNQTEVTVQLINYTKSGKKFWNVFHLQPMRDYKGDVQYFIGVQLDGTERLH
GAAEREAVCLIKKTAFQIAEAANDENYFGSGSGSVSKGEELFTGVVPILVELDGDVNGHKFSVS
GEGEGDATYGKLTLKLICTTGKLPVPWPTLVTTLGYGLQCFARYPDHMKQHDFFKSAMPEGYV
QERTIFFKDDGNYKTRAEVKFEGDTLVNRIELKGIDFKEDGNILGHKLEYNYNSHNVYITADKQK
NGIKANFKIRHNIEDGGVQLADHYQQNTPIGDGPVLLPDNHYLSYQSKLSKDPNEKRDHMVLLE
FVTAAGITLGMDELYKKKKKKKSKTKCVIM*

DNA Sequence:

ATGGTGTCTAAGGGCGAAGAGCTGATTAAGGAGAACATGCACATGAAGCTGTACATGGAGG
GCACCGTGAACAACCACCACTTCAAGTGCACATCCGAGGGCGAAGGCAAGCCCTACGAGG
GCACCCAGACCATGAGAATCAAGGTGGTCGAGGGCGGCCCTCTCCCCTTCGCCTTCGACA
TCCTGGCTACCAGCTTCATGTACGGCAGCAGAACCTTCATCAACCACACCCAGGGCATCCC
CGACTTCTTTAAGCAGTCCTTCCCTGAGGGCTTCACATGGGAGAGAGTCACCACATACGAA
GACGGGGGCGTGCTGACCGCTACCCAGGACACCAGCCTCCAGGACGGCTGCCTCATCTA
CAACGTCAAGATCAGAGGGGTGAACTTCCCATCCAACGGCCCTGTGATGCAGAAGAAAACA
CTCGGCTGGGAGGCCAACACCGAGATGCTGTACCCCGCTGACGGCGGCCTGGAAGGCAG
AACCGACATGGCCCTGAAGCTCGTGGGCGGGGGCCACCTGATCTGCAACTTCAAGACCAC
ATACAGATCCAAGAAACCCGCTAAGAACCTCAAGATGCCCGGCGTCTACTATGTGGACCAC
AGACTGGAAAGAATCAAGGAGGCCGACAAAGAGACCTACGTCGAGCAGCACGAGGTGGCT
GTGGCCAGATACTGCGACCTCCCTAGCAAACTGGGGCACAAACTTAATGGCATGGACGAGC
TGTACAAGTCCGGACTCAGATCTCGAGCTCAAGCTTCGAACGAATTCGGCATTGATCTGAG
CGGCCTGACCCTGCAGGAATTCAGCTCCCCGAAACGCCCTAAGCTGCTGCGTGAATATTAC
GATTGGCTGGTTGATAACAGCTTTACCCCATATCTGGTGGTGGATGCCACATACCTGGGCGT
GAACGTGCCCGTGGAGTATGTGAAAGACGGTCAGATCGTGCTGAATCTGTCTGCAAGTGC
GACCGGCAACCTGCAACTGACAAATGATTTTATCCAGTTCAACGCCCGCTTTAAGGGCGTG
TCTCGTGAACTGTATATCCCGATGGGTGCCGCTCTGGCCATTTACGCTCGCGAGAACGGCG
ATGGTGTGATGTTCGAACCAGAAGAAATCTATGACGAGCTGAATATTGGTCAGATCTCGTAC
GCCTCCCGGGGCGGTGGCTCATCTGGCGGAGGTGAGCTCCAGACAATTAAGTGTGTTGTT
GTGGGCGATGGTGCTTAA **---IRES Region---**
ATGGGGGTTGGTAAAACATGTCTCCTGATATCCTACACAACAAACAAATTTCCATCGGAGTAT
GTACCGACTGTTTTTGACAACTATGCAGTCACAGTTATGATTGGTGGAGAACCATATACTCTT

GGACTTTTTGATACTGCAGGGCTAGAGGATTATGACAGATTACGACCGCTGAGTTATCCACA
AACAGATGTATTTCTAGTCTGTTTTTCAGTGGTCTCTCCATCTTCATTTGAAAACGTGAAAGA
AAAGTGGGTGCCTGAGATAACTCACCACTGTCCAAAGACTCCTTTCTTGCTTGTTGGGACT
CAAATTGATCTCAGAGATGACCCCTCTACTATTGAGAAACTTGCCAAGAACAAACAGAAGCC
TATCACTCCAGAGACTGCTGAAAAGCTGGCCCGTGACCTGAAGGCTGTCAAGTATGTGGAG
TGTTCTGCACTTACACAGAGAGGTCTGAAGAATGTGTTTGATGAGGCTATCCTAGCTGCCCT
CGAGCCTCCGGAAACTCAACCCGGGGATCCCAATTGGGAGCTCGTGTACACGGCGCGCCT
GCAGGGAGGTGGCTCATCTGGCGGAGGTCAGATCTCGTACGCGTCCCGGGGCGAGTTTC
TGGCAACCACACTGGAACGGATCGAGAAAAATTTCGTGATTACTGATCCGAGACTGCCTGA
CAACCCAATCATTTTTGCGAGCGATTCCTTCCTGCAGCTGACAGAATATTCTCGGGAAGAGA
TCCTGGGGCGCAATTGCCGTTTTCTGCAGGGACCCGAGACAGACCGTGCCACTGTTCGGA
AAATCAGAGATGCTATTGACAACCAGACTGAAGTGACCGTTCAGCTGATCAATTATACCAAG
AGCGGCAAGAAGTTCTGGAACGTGTTCCACCTGCAGCCGATGCGCGATTATAAGGGCGAC
GTCCAGTACTTCATTGGCGTGCAGCTGGATGGCACCGAACGTCTTCATGGCGCCGCTGAG
CGTGAGGCGGTCTGCCTGATCAAAAAGACAGCCTTTCAGATTGCTGAGGCAGCGAACGAC
GAAAATTACTTTGGAAGCGGGAGTGGGAGCGTGAGCAAGGGCGAGGAGCTGTTCACCGG
GGTGGTGCCCATCCTGGTCGAGCTGGACGGCGACGTAAACGGCCACAAGTTCAGCGTGTC
CGGCGAGGGCGAGGGCGATGCCACCTACGGCAAGCTGACCCTGAAGCTCATCTGCACCA
CCGGCAAGCTGCCCGTGCCCTGGCCCACCCTCGTGACCACCCTCGGCTACGGCCTGCAG
TGCTTCGCCCGCTACCCCGACCACATGAAGCAGCACGACTTCTTCAAGTCCGCCATGCCC
GAAGGCTACGTCCAGGAGCGCACCATCTTCTTCAAGGACGACGGCAACTACAAGACCCGC
GCCGAGGTGAAGTTCGAGGGCGACACCCTGGTGAACCGCATCGAGCTGAAGGGCATCGA
CTTCAAGGAGGACGGCAACATCCTGGGGCACAAGCTGGAGTACAACTACAACAGCCACAA
CGTCTATATCACCGCCGACAAGCAGAAGAACGGCATCAAGGCCAACTTCAAGATCCGCCAC
AACATCGAGGACGGCGGCGTGCAGCTCGCCGACCACTACCAGCAGAACACCCCCATCGG
CGACGGCCCCGTGCTGCTGCCCGACAACCACTACCTGAGCTACCAGTCCAAGCTGAGCAA
AGACCCCAACGAGAAGCGCGATCACATGGTCCTGCTGGAGTTCGTGACCGCCGCCGGGAT
CACTCTCGGCATGGACGAGCTGTACAAGAAAAAAAAGAAGAAAAAGAGCAAGACCAAATGC
GTGATTATGTAA

**2) pIRES-tagRFPt-SspB Micro-Cdc42/N12-Cdc42/13C-iLID-mVenus-CAAX**

Amino Acid Sequence:

MVSKGEELIKENMHMKLYMEGTVNNHHFKCTSEGEGKPYEGTQTMRIKVVEGGPLPFAFDILA
TSFMYGSRTFINHTQGIPDFFKQSFPEGFTWERVTTYEDGGVLTATQDTSLQDGCLIYNVKIRG
VNFPSNGPVMQKKTLGWEANTEMLYPADGGLEGRTDMALKLVGGGHLICNFKTTYRSKKPAK
NLKMPGVYYVDHRLERIKEADKETYVEQHEVAVARYCDLPSKLGHKLNGMDELYKSGLRSRAQ
ASNEFGIDLSGLTLQEFSSPKRPKLLREYYDWLVDNSFTPYLVVDATYLGVNVPVEYVKDGQIVL
NLSASATGNLQLTNDFIQFNAQFKGVSRELYIPMGAALAIYARENGDGVMFEPEEIYDELNIGQIS
YASRGGGSSGGGELQTIKCVVVGDGA* **---IRES Region---**
MGVGKTCLLISYTTNKFPSEYVPTVFDNYAVTVMIGGEPYTLGLFDTAGLEDYDRLRPLSYPQT
DVFLVCFSVVSPSSFENVKEKWVPEITHHCPKTPFLLVGTQIDLRDDPSTIEKLAKNKQKPITPET
AEKLARDLKAVKYVECSALTQRGLKNVFDEAILAALEPPETQPGDPNWELVYTARLQGGGSSG
GGQISYASRGEFLATTLERIEKNFVITDPRLPDNPIIFASDSFLQLTEYSREEILGRNCRFLQGPET
DRATVRKIRDAIDNQTEVTVQLINYTKSGKKFWNVFHLQPMRDYKGDVQYFIGVQLDGTERLH
GAAEREAVCLIKKTAFQIAEAANDENYFGSGSGSVSKGEELFTGVVPILVELDGDVNGHKFSVS
GEGEGDATYGKLTLKLICTTGKLPVPWPTLVTTLGYGLQCFARYPDHMKQHDFFKSAMPEGYV
QERTIFFKDDGNYKTRAEVKFEGDTLVNRIELKGIDFKEDGNILGHKLEYNYNSHNVYITADKQK
NGIKANFKIRHNIEDGGVQLADHYQQNTPIGDGPVLLPDNHYLSYQSKLSKDPNEKRDHMVLLE
FVTAAGITLGMDELYKKKKKKKSKTKCVIM*

DNA Sequence:

ATGGTGTCTAAGGGCGAAGAGCTGATTAAGGAGAACATGCACATGAAGCTGTACATGGAGG
GCACCGTGAACAACCACCACTTCAAGTGCACATCCGAGGGCGAAGGCAAGCCCTACGAGG
GCACCCAGACCATGAGAATCAAGGTGGTCGAGGGCGGCCCTCTCCCCTTCGCCTTCGACA
TCCTGGCTACCAGCTTCATGTACGGCAGCAGAACCTTCATCAACCACACCCAGGGCATCCC
CGACTTCTTTAAGCAGTCCTTCCCTGAGGGCTTCACATGGGAGAGAGTCACCACATACGAA
GACGGGGGCGTGCTGACCGCTACCCAGGACACCAGCCTCCAGGACGGCTGCCTCATCTA
CAACGTCAAGATCAGAGGGGTGAACTTCCCATCCAACGGCCCTGTGATGCAGAAGAAAACA
CTCGGCTGGGAGGCCAACACCGAGATGCTGTACCCCGCTGACGGCGGCCTGGAAGGCAG
AACCGACATGGCCCTGAAGCTCGTGGGCGGGGGCCACCTGATCTGCAACTTCAAGACCAC
ATACAGATCCAAGAAACCCGCTAAGAACCTCAAGATGCCCGGCGTCTACTATGTGGACCAC
AGACTGGAAAGAATCAAGGAGGCCGACAAAGAGACCTACGTCGAGCAGCACGAGGTGGCT
GTGGCCAGATACTGCGACCTCCCTAGCAAACTGGGGCACAAACTTAATGGCATGGACGAGC
TGTACAAGTCCGGACTCAGATCTCGAGCTCAAGCTTCGAACGAATTCGGCATTGATCTGAG
CGGCCTGACCCTGCAGGAATTCAGCTCCCCGAAACGCCCTAAGCTGCTGCGTGAATATTAC
GATTGGCTGGTTGATAACAGCTTTACCCCATATCTGGTGGTGGATGCCACATACCTGGGCGT
GAACGTGCCCGTGGAGTATGTGAAAGACGGTCAGATCGTGCTGAATCTGTCTGCAAGTGC
GACCGGCAACCTGCAACTGACAAATGATTTTATCCAGTTCAACGCCCAGTTTAAGGGCGTG
TCTCGTGAACTGTATATCCCGATGGGTGCCGCTCTGGCCATTTACGCTCGCGAGAACGGCG
ATGGTGTGATGTTCGAACCAGAAGAAATCTATGACGAGCTGAATATTGGTCAGATCTCGTAC
GCCTCCCGGGGCGGTGGCTCATCTGGCGGAGGTGAGCTCCAGACAATTAAGTGTGTTGTT
GTGGGCGATGGTGCTTAA **---IRES Region---**
ATGGGGGTTGGTAAAACATGTCTCCTGATATCCTACACAACAAACAAATTTCCATCGGAGTAT
GTACCGACTGTTTTTGACAACTATGCAGTCACAGTTATGATTGGTGGAGAACCATATACTCTT
GGACTTTTTGATACTGCAGGGCTAGAGGATTATGACAGATTACGACCGCTGAGTTATCCACA
AACAGATGTATTTCTAGTCTGTTTTTCAGTGGTCTCTCCATCTTCATTTGAAAACGTGAAAGA
AAAGTGGGTGCCTGAGATAACTCACCACTGTCCAAAGACTCCTTTCTTGCTTGTTGGGACT
CAAATTGATCTCAGAGATGACCCCTCTACTATTGAGAAACTTGCCAAGAACAAACAGAAGCC

TATCACTCCAGAGACTGCTGAAAAGCTGGCCCGTGACCTGAAGGCTGTCAAGTATGTGGAG
TGTTCTGCACTTACACAGAGAGGTCTGAAGAATGTGTTTGATGAGGCTATCCTAGCTGCCCT
CGAGCCTCCGGAAACTCAACCCGGGGATCCCAATTGGGAGCTCGTGTACACGGCGCGCCT
GCAGGGAGGTGGCTCATCTGGCGGAGGTCAGATCTCGTACGCGTCCCGGGGCGAGTTTC
TGGCAACCACACTGGAACGGATCGAGAAAAATTTCGTGATTACTGATCCGAGACTGCCTGA
CAACCCAATCATTTTTGCGAGCGATTCCTTCCTGCAGCTGACAGAATATTCTCGGGAAGAGA
TCCTGGGGCGCAATTGCCGTTTTCTGCAGGGACCCGAGACAGACCGTGCCACTGTTCGGA
AAATCAGAGATGCTATTGACAACCAGACTGAAGTGACCGTTCAGCTGATCAATTATACCAAG
AGCGGCAAGAAGTTCTGGAACGTGTTCCACCTGCAGCCGATGCGCGATTATAAGGGCGAC
GTCCAGTACTTCATTGGCGTGCAGCTGGATGGCACCGAACGTCTTCATGGCGCCGCTGAG
CGTGAGGCGGTCTGCCTGATCAAAAAGACAGCCTTTCAGATTGCTGAGGCAGCGAACGAC
GAAAATTACTTTGGAAGCGGGAGTGGGAGCGTGAGCAAGGGCGAGGAGCTGTTCACCGG
GGTGGTGCCCATCCTGGTCGAGCTGGACGGCGACGTAAACGGCCACAAGTTCAGCGTGTC
CGGCGAGGGCGAGGGCGATGCCACCTACGGCAAGCTGACCCTGAAGCTCATCTGCACCA
CCGGCAAGCTGCCCGTGCCCTGGCCCACCCTCGTGACCACCCTCGGCTACGGCCTGCAG
TGCTTCGCCCGCTACCCCGACCACATGAAGCAGCACGACTTCTTCAAGTCCGCCATGCCC
GAAGGCTACGTCCAGGAGCGCACCATCTTCTTCAAGGACGACGGCAACTACAAGACCCGC
GCCGAGGTGAAGTTCGAGGGCGACACCCTGGTGAACCGCATCGAGCTGAAGGGCATCGA
CTTCAAGGAGGACGGCAACATCCTGGGGCACAAGCTGGAGTACAACTACAACAGCCACAA
CGTCTATATCACCGCCGACAAGCAGAAGAACGGCATCAAGGCCAACTTCAAGATCCGCCAC
AACATCGAGGACGGCGGCGTGCAGCTCGCCGACCACTACCAGCAGAACACCCCCATCGG
CGACGGCCCCGTGCTGCTGCCCGACAACCACTACCTGAGCTACCAGTCCAAGCTGAGCAA
AGACCCCAACGAGAAGCGCGATCACATGGTCCTGCTGGAGTTCGTGACCGCCGCCGGGAT
CACTCTCGGCATGGACGAGCTGTACAAGAAAAAAAAGAAGAAAAAGAGCAAGACCAAATGC
GTGATTATGTAA

#### 3) pIRES-tagRFPt-SspB Milli-Cdc42/N12-Cdc42/13C-iLID-mVenus-CAAX

Amino Acid Sequence:

MVSKGEELIKENMHMKLYMEGTVNNHHFKCTSEGEGKPYEGTQTMRIKVVEGGPLPFAFDILA
TSFMYGSRTFINHTQGIPDFFKQSFPEGFTWERVTTYEDGGVLTATQDTSLQDGCLIYNVKIRG
VNFPSNGPVMQKKTLGWEANTEMLYPADGGLEGRTDMALKLVGGGHLICNFKTTYRSKKPAK
NLKMPGVYYVDHRLERIKEADKETYVEQHEVAVARYCDLPSKLGHKLNGMDELYKSGLRSRAQ
ASNEFGIDLSGLTLQEFSSPKRPKLLREYYDWLVDNSFTPYLVVDATYLGVNVPVEYVKDGQIVL
NLSASVTGNLQLTNDFIQFNAQFKGVSRELYIPMGAALAIYARENGDGVMFEPEEIYDELNIGQIS
YASRGGGSSGGGELQTIKCVVVGDGA* **---IRES Region---**
MGVGKTCLLISYTTNKFPSEYVPTVFDNYAVTVMIGGEPYTLGLFDTAGLEDYDRLRPLSYPQT
DVFLVCFSVVSPSSFENVKEKWVPEITHHCPKTPFLLVGTQIDLRDDPSTIEKLAKNKQKPITPET
AEKLARDLKAVKYVECSALTQRGLKNVFDEAILAALEPPETQPGDPNWELVYTARLQGGGSSG
GGQISYASRGEFLATTLERIEKNFVITDPRLPDNPIIFASDSFLQLTEYSREEILGRNCRFLQGPET
DRATVRKIRDAIDNQTEVTVQLINYTKSGKKFWNVFHLQPMRDYKGDVQYFIGVQLDGTERLH
GAAEREAVCLIKKTAFQIAEAANDENYFGSGSGSVSKGEELFTGVVPILVELDGDVNGHKFSVS
GEGEGDATYGKLTLKLICTTGKLPVPWPTLVTTLGYGLQCFARYPDHMKQHDFFKSAMPEGYV
QERTIFFKDDGNYKTRAEVKFEGDTLVNRIELKGIDFKEDGNILGHKLEYNYNSHNVYITADKQK
NGIKANFKIRHNIEDGGVQLADHYQQNTPIGDGPVLLPDNHYLSYQSKLSKDPNEKRDHMVLLE
FVTAAGITLGMDELYKKKKKKKSKTKCVIM*

DNA Sequence:

ATGGTGTCTAAGGGCGAAGAGCTGATTAAGGAGAACATGCACATGAAGCTGTACATGGAGG
GCACCGTGAACAACCACCACTTCAAGTGCACATCCGAGGGCGAAGGCAAGCCCTACGAGG
GCACCCAGACCATGAGAATCAAGGTGGTCGAGGGCGGCCCTCTCCCCTTCGCCTTCGACA
TCCTGGCTACCAGCTTCATGTACGGCAGCAGAACCTTCATCAACCACACCCAGGGCATCCC
CGACTTCTTTAAGCAGTCCTTCCCTGAGGGCTTCACATGGGAGAGAGTCACCACATACGAA
GACGGGGGCGTGCTGACCGCTACCCAGGACACCAGCCTCCAGGACGGCTGCCTCATCTA
CAACGTCAAGATCAGAGGGGTGAACTTCCCATCCAACGGCCCTGTGATGCAGAAGAAAACA
CTCGGCTGGGAGGCCAACACCGAGATGCTGTACCCCGCTGACGGCGGCCTGGAAGGCAG
AACCGACATGGCCCTGAAGCTCGTGGGCGGGGGCCACCTGATCTGCAACTTCAAGACCAC
ATACAGATCCAAGAAACCCGCTAAGAACCTCAAGATGCCCGGCGTCTACTATGTGGACCAC
AGACTGGAAAGAATCAAGGAGGCCGACAAAGAGACCTACGTCGAGCAGCACGAGGTGGCT
GTGGCCAGATACTGCGACCTCCCTAGCAAACTGGGGCACAAACTTAATGGCATGGACGAGC
TGTACAAGTCCGGACTCAGATCTCGAGCTCAAGCTTCGAACGAATTCGGCATTGATCTGAG
CGGCCTGACCCTGCAGGAATTCAGCTCCCCGAAACGCCCTAAGCTGCTGCGTGAATATTAC
GATTGGCTGGTTGATAACAGCTTTACCCCATATCTGGTGGTGGATGCCACATACCTGGGCGT
GAACGTGCCCGTGGAGTATGTGAAAGACGGTCAGATCGTGCTGAATCTGTCTGCAAGTGTG
ACCGGCAACCTGCAACTGACAAATGATTTTATCCAGTTCAACGCCCAGTTTAAGGGCGTGTC
TCGTGAACTGTATATCCCGATGGGTGCCGCTCTGGCCATTTACGCTCGCGAGAACGGCGAT
GGTGTGATGTTCGAACCAGAAGAAATCTATGACGAGCTGAATATTGGTCAGATCTCGTACGC
CTCCCGGGGCGGTGGCTCATCTGGCGGAGGTGAGCTCCAGACAATTAAGTGTGTTGTTGT
GGGCGATGGTGCTTAA **---IRES Region---**
ATGGGGGTTGGTAAAACATGTCTCCTGATATCCTACACAACAAACAAATTTCCATCGGAGTAT
GTACCGACTGTTTTTGACAACTATGCAGTCACAGTTATGATTGGTGGAGAACCATATACTCTT
GGACTTTTTGATACTGCAGGGCTAGAGGATTATGACAGATTACGACCGCTGAGTTATCCACA
AACAGATGTATTTCTAGTCTGTTTTTCAGTGGTCTCTCCATCTTCATTTGAAAACGTGAAAGA
AAAGTGGGTGCCTGAGATAACTCACCACTGTCCAAAGACTCCTTTCTTGCTTGTTGGGACT
CAAATTGATCTCAGAGATGACCCCTCTACTATTGAGAAACTTGCCAAGAACAAACAGAAGCC

TATCACTCCAGAGACTGCTGAAAAGCTGGCCCGTGACCTGAAGGCTGTCAAGTATGTGGAG
TGTTCTGCACTTACACAGAGAGGTCTGAAGAATGTGTTTGATGAGGCTATCCTAGCTGCCCT
CGAGCCTCCGGAAACTCAACCCGGGGATCCCAATTGGGAGCTCGTGTACACGGCGCGCCT
GCAGGGAGGTGGCTCATCTGGCGGAGGTCAGATCTCGTACGCGTCCCGGGGCGAGTTTC
TGGCAACCACACTGGAACGGATCGAGAAAAATTTCGTGATTACTGATCCGAGACTGCCTGA
CAACCCAATCATTTTTGCGAGCGATTCCTTCCTGCAGCTGACAGAATATTCTCGGGAAGAGA
TCCTGGGGCGCAATTGCCGTTTTCTGCAGGGACCCGAGACAGACCGTGCCACTGTTCGGA
AAATCAGAGATGCTATTGACAACCAGACTGAAGTGACCGTTCAGCTGATCAATTATACCAAG
AGCGGCAAGAAGTTCTGGAACGTGTTCCACCTGCAGCCGATGCGCGATTATAAGGGCGAC
GTCCAGTACTTCATTGGCGTGCAGCTGGATGGCACCGAACGTCTTCATGGCGCCGCTGAG
CGTGAGGCGGTCTGCCTGATCAAAAAGACAGCCTTTCAGATTGCTGAGGCAGCGAACGAC
GAAAATTACTTTGGAAGCGGGAGTGGGAGCGTGAGCAAGGGCGAGGAGCTGTTCACCGG
GGTGGTGCCCATCCTGGTCGAGCTGGACGGCGACGTAAACGGCCACAAGTTCAGCGTGTC
CGGCGAGGGCGAGGGCGATGCCACCTACGGCAAGCTGACCCTGAAGCTCATCTGCACCA
CCGGCAAGCTGCCCGTGCCCTGGCCCACCCTCGTGACCACCCTCGGCTACGGCCTGCAG
TGCTTCGCCCGCTACCCCGACCACATGAAGCAGCACGACTTCTTCAAGTCCGCCATGCCC
GAAGGCTACGTCCAGGAGCGCACCATCTTCTTCAAGGACGACGGCAACTACAAGACCCGC
GCCGAGGTGAAGTTCGAGGGCGACACCCTGGTGAACCGCATCGAGCTGAAGGGCATCGA
CTTCAAGGAGGACGGCAACATCCTGGGGCACAAGCTGGAGTACAACTACAACAGCCACAA
CGTCTATATCACCGCCGACAAGCAGAAGAACGGCATCAAGGCCAACTTCAAGATCCGCCAC
AACATCGAGGACGGCGGCGTGCAGCTCGCCGACCACTACCAGCAGAACACCCCCATCGG
CGACGGCCCCGTGCTGCTGCCCGACAACCACTACCTGAGCTACCAGTCCAAGCTGAGCAA
AGACCCCAACGAGAAGCGCGATCACATGGTCCTGCTGGAGTTCGTGACCGCCGCCGGGAT
CACTCTCGGCATGGACGAGCTGTACAAGAAAAAAAAGAAGAAAAAGAGCAAGACCAAATGC
GTGATTATGTAA

#### 4) pIRES-tagRFPt-SspB Nano-Cdc42/N12-Cdc42/13C-iLID C252A-mVenus-CAAX

Amino Acid Sequence (C252A is underlined):

MVSKGEELIKENMHMKLYMEGTVNNHHFKCTSEGEGKPYEGTQTMRIKVVEGGPLPFAFDILA
TSFMYGSRTFINHTQGIPDFFKQSFPEGFTWERVTTYEDGGVLTATQDTSLQDGCLIYNVKIRG
VNFPSNGPVMQKKTLGWEANTEMLYPADGGLEGRTDMALKLVGGGHLICNFKTTYRSKKPAK
NLKMPGVYYVDHRLERIKEADKETYVEQHEVAVARYCDLPSKLGHKLNGMDELYKSGLRSRAQ
ASNEFGIDLSGLTLQEFSSPKRPKLLREYYDWLVDNSFTPYLVVDATYLGVNVPVEYVKDGQIVL
NLSASATGNLQLTNDFIQFNARFKGVSRELYIPMGAALAIYARENGDGVMFEPEEIYDELNIGQIS
YASRGGGSSGGGELQTIKCVVVGDGA* **---IRES Region---**
MGVGKTCLLISYTTNKFPSEYVPTVFDNYAVTVMIGGEPYTLGLFDTAGLEDYDRLRPLSYPQT
DVFLVCFSVVSPSSFENVKEKWVPEITHHCPKTPFLLVGTQIDLRDDPSTIEKLAKNKQKPITPET
AEKLARDLKAVKYVECSALTQRGLKNVFDEAILAALEPPETQPGDPNWELVYTARLQGGGSSG
GGQISYASRGEFLATTLERIEKNFVITDPRLPDNPIIFASDSFLQLTEYSREEILGRN**A**RFLQGPET
DRATVRKIRDAIDNQTEVTVQLINYTKSGKKFWNVFHLQPMRDYKGDVQYFIGVQLDGTERLH
GAAEREAVCLIKKTAFQIAEAANDENYFGSGSGSVSKGEELFTGVVPILVELDGDVNGHKFSVS
GEGEGDATYGKLTLKLICTTGKLPVPWPTLVTTLGYGLQCFARYPDHMKQHDFFKSAMPEGYV
QERTIFFKDDGNYKTRAEVKFEGDTLVNRIELKGIDFKEDGNILGHKLEYNYNSHNVYITADKQK
NGIKANFKIRHNIEDGGVQLADHYQQNTPIGDGPVLLPDNHYLSYQSKLSKDPNEKRDHMVLLE
FVTAAGITLGMDELYKKKKKKKSKTKCVIM*

DNA Sequence:

ATGGTGTCTAAGGGCGAAGAGCTGATTAAGGAGAACATGCACATGAAGCTGTACATGGAGG
GCACCGTGAACAACCACCACTTCAAGTGCACATCCGAGGGCGAAGGCAAGCCCTACGAGG
GCACCCAGACCATGAGAATCAAGGTGGTCGAGGGCGGCCCTCTCCCCTTCGCCTTCGACA
TCCTGGCTACCAGCTTCATGTACGGCAGCAGAACCTTCATCAACCACACCCAGGGCATCCC
CGACTTCTTTAAGCAGTCCTTCCCTGAGGGCTTCACATGGGAGAGAGTCACCACATACGAA
GACGGGGGCGTGCTGACCGCTACCCAGGACACCAGCCTCCAGGACGGCTGCCTCATCTA
CAACGTCAAGATCAGAGGGGTGAACTTCCCATCCAACGGCCCTGTGATGCAGAAGAAAACA
CTCGGCTGGGAGGCCAACACCGAGATGCTGTACCCCGCTGACGGCGGCCTGGAAGGCAG
AACCGACATGGCCCTGAAGCTCGTGGGCGGGGGCCACCTGATCTGCAACTTCAAGACCAC
ATACAGATCCAAGAAACCCGCTAAGAACCTCAAGATGCCCGGCGTCTACTATGTGGACCAC
AGACTGGAAAGAATCAAGGAGGCCGACAAAGAGACCTACGTCGAGCAGCACGAGGTGGCT
GTGGCCAGATACTGCGACCTCCCTAGCAAACTGGGGCACAAACTTAATGGCATGGACGAGC
TGTACAAGTCCGGACTCAGATCTCGAGCTCAAGCTTCGAACGAATTCGGCATTGATCTGAG
CGGCCTGACCCTGCAGGAATTCAGCTCCCCGAAACGCCCTAAGCTGCTGCGTGAATATTAC
GATTGGCTGGTTGATAACAGCTTTACCCCATATCTGGTGGTGGATGCCACATACCTGGGCGT
GAACGTGCCCGTGGAGTATGTGAAAGACGGTCAGATCGTGCTGAATCTGTCTGCAAGTGC
GACCGGCAACCTGCAACTGACAAATGATTTTATCCAGTTCAACGCCCGCTTTAAGGGCGTG
TCTCGTGAACTGTATATCCCGATGGGTGCCGCTCTGGCCATTTACGCTCGCGAGAACGGCG
ATGGTGTGATGTTCGAACCAGAAGAAATCTATGACGAGCTGAATATTGGTCAGATCTCGTAC
GCCTCCCGGGGCGGTGGCTCATCTGGCGGAGGTGAGCTCCAGACAATTAAGTGTGTTGTT
GTGGGCGATGGTGCTTAA **---IRES Region---**
ATGGGGGTTGGTAAAACATGTCTCCTGATATCCTACACAACAAACAAATTTCCATCGGAGTAT
GTACCGACTGTTTTTGACAACTATGCAGTCACAGTTATGATTGGTGGAGAACCATATACTCTT
GGACTTTTTGATACTGCAGGGCTAGAGGATTATGACAGATTACGACCGCTGAGTTATCCACA
AACAGATGTATTTCTAGTCTGTTTTTCAGTGGTCTCTCCATCTTCATTTGAAAACGTGAAAGA
AAAGTGGGTGCCTGAGATAACTCACCACTGTCCAAAGACTCCTTTCTTGCTTGTTGGGACT
CAAATTGATCTCAGAGATGACCCCTCTACTATTGAGAAACTTGCCAAGAACAAACAGAAGCC

TATCACTCCAGAGACTGCTGAAAAGCTGGCCCGTGACCTGAAGGCTGTCAAGTATGTGGAG
TGTTCTGCACTTACACAGAGAGGTCTGAAGAATGTGTTTGATGAGGCTATCCTAGCTGCCCT
CGAGCCTCCGGAAACTCAACCCGGGGATCCCAATTGGGAGCTCGTGTACACGGCGCGCCT
GCAGGGAGGTGGCTCATCTGGCGGAGGTCAGATCTCGTACGCGTCCCGGGGCGAGTTTC
TGGCAACCACACTGGAACGGATCGAGAAAAATTTCGTGATTACTGATCCGAGACTGCCTGA
CAACCCAATCATTTTTGCGAGCGATTCCTTCCTGCAGCTGACAGAATATTCTCGGGAAGAGA
TCCTGGGGCGCAATGCCCGTTTTCTGCAGGGACCCGAGACAGACCGTGCCACTGTTCGGA
AAATCAGAGATGCTATTGACAACCAGACTGAAGTGACCGTTCAGCTGATCAATTATACCAAG
AGCGGCAAGAAGTTCTGGAACGTGTTCCACCTGCAGCCGATGCGCGATTATAAGGGCGAC
GTCCAGTACTTCATTGGCGTGCAGCTGGATGGCACCGAACGTCTTCATGGCGCCGCTGAG
CGTGAGGCGGTCTGCCTGATCAAAAAGACAGCCTTTCAGATTGCTGAGGCAGCGAACGAC
GAAAATTACTTTGGAAGCGGGAGTGGGAGCGTGAGCAAGGGCGAGGAGCTGTTCACCGG
GGTGGTGCCCATCCTGGTCGAGCTGGACGGCGACGTAAACGGCCACAAGTTCAGCGTGTC
CGGCGAGGGCGAGGGCGATGCCACCTACGGCAAGCTGACCCTGAAGCTCATCTGCACCA
CCGGCAAGCTGCCCGTGCCCTGGCCCACCCTCGTGACCACCCTCGGCTACGGCCTGCAG
TGCTTCGCCCGCTACCCCGACCACATGAAGCAGCACGACTTCTTCAAGTCCGCCATGCCC
GAAGGCTACGTCCAGGAGCGCACCATCTTCTTCAAGGACGACGGCAACTACAAGACCCGC
GCCGAGGTGAAGTTCGAGGGCGACACCCTGGTGAACCGCATCGAGCTGAAGGGCATCGA
CTTCAAGGAGGACGGCAACATCCTGGGGCACAAGCTGGAGTACAACTACAACAGCCACAA
CGTCTATATCACCGCCGACAAGCAGAAGAACGGCATCAAGGCCAACTTCAAGATCCGCCAC
AACATCGAGGACGGCGGCGTGCAGCTCGCCGACCACTACCAGCAGAACACCCCCATCGG
CGACGGCCCCGTGCTGCTGCCCGACAACCACTACCTGAGCTACCAGTCCAAGCTGAGCAA
AGACCCCAACGAGAAGCGCGATCACATGGTCCTGCTGGAGTTCGTGACCGCCGCCGGGAT
CACTCTCGGCATGGACGAGCTGTACAAGAAAAAAAAGAAGAAAAAGAGCAAGACCAAATGC
GTGATTATGTAA

#### 5) pIRES-tagRFPt-SspB Micro-Cdc42/N12-Cdc42/13C-iLID C252A-mVenus-CAAX

Amino Acid Sequence (C252A is underlined):

MVSKGEELIKENMHMKLYMEGTVNNHHFKCTSEGEGKPYEGTQTMRIKVVEGGPLPFAFDILA
TSFMYGSRTFINHTQGIPDFFKQSFPEGFTWERVTTYEDGGVLTATQDTSLQDGCLIYNVKIRG
VNFPSNGPVMQKKTLGWEANTEMLYPADGGLEGRTDMALKLVGGGHLICNFKTTYRSKKPAK
NLKMPGVYYVDHRLERIKEADKETYVEQHEVAVARYCDLPSKLGHKLNGMDELYKSGLRSRAQ
ASNEFGIDLSGLTLQEFSSPKRPKLLREYYDWLVDNSFTPYLVVDATYLGVNVPVEYVKDGQIVL
NLSASATGNLQLTNDFIQFNAQFKGVSRELYIPMGAALAIYARENGDGVMFEPEEIYDELNIGQIS
YASRGGGSSGGGELQTIKCVVVGDGA* **---IRES Region---**
MGVGKTCLLISYTTNKFPSEYVPTVFDNYAVTVMIGGEPYTLGLFDTAGLEDYDRLRPLSYPQT
DVFLVCFSVVSPSSFENVKEKWVPEITHHCPKTPFLLVGTQIDLRDDPSTIEKLAKNKQKPITPET
AEKLARDLKAVKYVECSALTQRGLKNVFDEAILAALEPPETQPGDPNWELVYTARLQGGGSSG
GGQISYASRGEFLATTLERIEKNFVITDPRLPDNPIIFASDSFLQLTEYSREEILGRN**A**RFLQGPET
DRATVRKIRDAIDNQTEVTVQLINYTKSGKKFWNVFHLQPMRDYKGDVQYFIGVQLDGTERLH
GAAEREAVCLIKKTAFQIAEAANDENYFGSGSGSVSKGEELFTGVVPILVELDGDVNGHKFSVS
GEGEGDATYGKLTLKLICTTGKLPVPWPTLVTTLGYGLQCFARYPDHMKQHDFFKSAMPEGYV
QERTIFFKDDGNYKTRAEVKFEGDTLVNRIELKGIDFKEDGNILGHKLEYNYNSHNVYITADKQK
NGIKANFKIRHNIEDGGVQLADHYQQNTPIGDGPVLLPDNHYLSYQSKLSKDPNEKRDHMVLLE
FVTAAGITLGMDELYKKKKKKKSKTKCVIM*

DNA Sequence:

ATGGTGTCTAAGGGCGAAGAGCTGATTAAGGAGAACATGCACATGAAGCTGTACATGGAGG
GCACCGTGAACAACCACCACTTCAAGTGCACATCCGAGGGCGAAGGCAAGCCCTACGAGG
GCACCCAGACCATGAGAATCAAGGTGGTCGAGGGCGGCCCTCTCCCCTTCGCCTTCGACA
TCCTGGCTACCAGCTTCATGTACGGCAGCAGAACCTTCATCAACCACACCCAGGGCATCCC
CGACTTCTTTAAGCAGTCCTTCCCTGAGGGCTTCACATGGGAGAGAGTCACCACATACGAA
GACGGGGGCGTGCTGACCGCTACCCAGGACACCAGCCTCCAGGACGGCTGCCTCATCTA
CAACGTCAAGATCAGAGGGGTGAACTTCCCATCCAACGGCCCTGTGATGCAGAAGAAAACA
CTCGGCTGGGAGGCCAACACCGAGATGCTGTACCCCGCTGACGGCGGCCTGGAAGGCAG
AACCGACATGGCCCTGAAGCTCGTGGGCGGGGGCCACCTGATCTGCAACTTCAAGACCAC
ATACAGATCCAAGAAACCCGCTAAGAACCTCAAGATGCCCGGCGTCTACTATGTGGACCAC
AGACTGGAAAGAATCAAGGAGGCCGACAAAGAGACCTACGTCGAGCAGCACGAGGTGGCT
GTGGCCAGATACTGCGACCTCCCTAGCAAACTGGGGCACAAACTTAATGGCATGGACGAGC
TGTACAAGTCCGGACTCAGATCTCGAGCTCAAGCTTCGAACGAATTCGGCATTGATCTGAG
CGGCCTGACCCTGCAGGAATTCAGCTCCCCGAAACGCCCTAAGCTGCTGCGTGAATATTAC
GATTGGCTGGTTGATAACAGCTTTACCCCATATCTGGTGGTGGATGCCACATACCTGGGCGT
GAACGTGCCCGTGGAGTATGTGAAAGACGGTCAGATCGTGCTGAATCTGTCTGCAAGTGC
GACCGGCAACCTGCAACTGACAAATGATTTTATCCAGTTCAACGCCCAGTTTAAGGGCGTG
TCTCGTGAACTGTATATCCCGATGGGTGCCGCTCTGGCCATTTACGCTCGCGAGAACGGCG
ATGGTGTGATGTTCGAACCAGAAGAAATCTATGACGAGCTGAATATTGGTCAGATCTCGTAC
GCCTCCCGGGGCGGTGGCTCATCTGGCGGAGGTGAGCTCCAGACAATTAAGTGTGTTGTT
GTGGGCGATGGTGCTTAA **---IRES Region---**
ATGGGGGTTGGTAAAACATGTCTCCTGATATCCTACACAACAAACAAATTTCCATCGGAGTAT
GTACCGACTGTTTTTGACAACTATGCAGTCACAGTTATGATTGGTGGAGAACCATATACTCTT
GGACTTTTTGATACTGCAGGGCTAGAGGATTATGACAGATTACGACCGCTGAGTTATCCACA
AACAGATGTATTTCTAGTCTGTTTTTCAGTGGTCTCTCCATCTTCATTTGAAAACGTGAAAGA
AAAGTGGGTGCCTGAGATAACTCACCACTGTCCAAAGACTCCTTTCTTGCTTGTTGGGACT
CAAATTGATCTCAGAGATGACCCCTCTACTATTGAGAAACTTGCCAAGAACAAACAGAAGCC

TATCACTCCAGAGACTGCTGAAAAGCTGGCCCGTGACCTGAAGGCTGTCAAGTATGTGGAG
TGTTCTGCACTTACACAGAGAGGTCTGAAGAATGTGTTTGATGAGGCTATCCTAGCTGCCCT
CGAGCCTCCGGAAACTCAACCCGGGGATCCCAATTGGGAGCTCGTGTACACGGCGCGCCT
GCAGGGAGGTGGCTCATCTGGCGGAGGTCAGATCTCGTACGCGTCCCGGGGCGAGTTTC
TGGCAACCACACTGGAACGGATCGAGAAAAATTTCGTGATTACTGATCCGAGACTGCCTGA
CAACCCAATCATTTTTGCGAGCGATTCCTTCCTGCAGCTGACAGAATATTCTCGGGAAGAGA
TCCTGGGGCGCAATGCCCGTTTTCTGCAGGGACCCGAGACAGACCGTGCCACTGTTCGGA
AAATCAGAGATGCTATTGACAACCAGACTGAAGTGACCGTTCAGCTGATCAATTATACCAAG
AGCGGCAAGAAGTTCTGGAACGTGTTCCACCTGCAGCCGATGCGCGATTATAAGGGCGAC
GTCCAGTACTTCATTGGCGTGCAGCTGGATGGCACCGAACGTCTTCATGGCGCCGCTGAG
CGTGAGGCGGTCTGCCTGATCAAAAAGACAGCCTTTCAGATTGCTGAGGCAGCGAACGAC
GAAAATTACTTTGGAAGCGGGAGTGGGAGCGTGAGCAAGGGCGAGGAGCTGTTCACCGG
GGTGGTGCCCATCCTGGTCGAGCTGGACGGCGACGTAAACGGCCACAAGTTCAGCGTGTC
CGGCGAGGGCGAGGGCGATGCCACCTACGGCAAGCTGACCCTGAAGCTCATCTGCACCA
CCGGCAAGCTGCCCGTGCCCTGGCCCACCCTCGTGACCACCCTCGGCTACGGCCTGCAG
TGCTTCGCCCGCTACCCCGACCACATGAAGCAGCACGACTTCTTCAAGTCCGCCATGCCC
GAAGGCTACGTCCAGGAGCGCACCATCTTCTTCAAGGACGACGGCAACTACAAGACCCGC
GCCGAGGTGAAGTTCGAGGGCGACACCCTGGTGAACCGCATCGAGCTGAAGGGCATCGA
CTTCAAGGAGGACGGCAACATCCTGGGGCACAAGCTGGAGTACAACTACAACAGCCACAA
CGTCTATATCACCGCCGACAAGCAGAAGAACGGCATCAAGGCCAACTTCAAGATCCGCCAC
AACATCGAGGACGGCGGCGTGCAGCTCGCCGACCACTACCAGCAGAACACCCCCATCGG
CGACGGCCCCGTGCTGCTGCCCGACAACCACTACCTGAGCTACCAGTCCAAGCTGAGCAA
AGACCCCAACGAGAAGCGCGATCACATGGTCCTGCTGGAGTTCGTGACCGCCGCCGGGAT
CACTCTCGGCATGGACGAGCTGTACAAGAAAAAAAAGAAGAAAAAGAGCAAGACCAAATGC
GTGATTATGTAA

#### 6) pIRES-tagRFPt-SspB Milli-Cdc42/N12-Cdc42/13C-iLID C252A-mVenus-CAAX

Amino Acid Sequence (C252A is underlined):

MVSKGEELIKENMHMKLYMEGTVNNHHFKCTSEGEGKPYEGTQTMRIKVVEGGPLPFAFDILA
TSFMYGSRTFINHTQGIPDFFKQSFPEGFTWERVTTYEDGGVLTATQDTSLQDGCLIYNVKIRG
VNFPSNGPVMQKKTLGWEANTEMLYPADGGLEGRTDMALKLVGGGHLICNFKTTYRSKKPAK
NLKMPGVYYVDHRLERIKEADKETYVEQHEVAVARYCDLPSKLGHKLNGMDELYKSGLRSRAQ
ASNEFGIDLSGLTLQEFSSPKRPKLLREYYDWLVDNSFTPYLVVDATYLGVNVPVEYVKDGQIVL
NLSASVTGNLQLTNDFIQFNAQFKGVSRELYIPMGAALAIYARENGDGVMFEPEEIYDELNIGQIS
YASRGGGSSGGGELQTIKCVVVGDGA* **---IRES Region---**
MGVGKTCLLISYTTNKFPSEYVPTVFDNYAVTVMIGGEPYTLGLFDTAGLEDYDRLRPLSYPQT
DVFLVCFSVVSPSSFENVKEKWVPEITHHCPKTPFLLVGTQIDLRDDPSTIEKLAKNKQKPITPET
AEKLARDLKAVKYVECSALTQRGLKNVFDEAILAALEPPETQPGDPNWELVYTARLQGGGSSG
GGQISYASRGEFLATTLERIEKNFVITDPRLPDNPIIFASDSFLQLTEYSREEILGRN**A**RFLQGPET
DRATVRKIRDAIDNQTEVTVQLINYTKSGKKFWNVFHLQPMRDYKGDVQYFIGVQLDGTERLH
GAAEREAVCLIKKTAFQIAEAANDENYFGSGSGSVSKGEELFTGVVPILVELDGDVNGHKFSVS
GEGEGDATYGKLTLKLICTTGKLPVPWPTLVTTLGYGLQCFARYPDHMKQHDFFKSAMPEGYV
QERTIFFKDDGNYKTRAEVKFEGDTLVNRIELKGIDFKEDGNILGHKLEYNYNSHNVYITADKQK
NGIKANFKIRHNIEDGGVQLADHYQQNTPIGDGPVLLPDNHYLSYQSKLSKDPNEKRDHMVLLE
FVTAAGITLGMDELYKKKKKKKSKTKCVIM*

DNA Sequence:

ATGGTGTCTAAGGGCGAAGAGCTGATTAAGGAGAACATGCACATGAAGCTGTACATGGAGG
GCACCGTGAACAACCACCACTTCAAGTGCACATCCGAGGGCGAAGGCAAGCCCTACGAGG
GCACCCAGACCATGAGAATCAAGGTGGTCGAGGGCGGCCCTCTCCCCTTCGCCTTCGACA
TCCTGGCTACCAGCTTCATGTACGGCAGCAGAACCTTCATCAACCACACCCAGGGCATCCC
CGACTTCTTTAAGCAGTCCTTCCCTGAGGGCTTCACATGGGAGAGAGTCACCACATACGAA
GACGGGGGCGTGCTGACCGCTACCCAGGACACCAGCCTCCAGGACGGCTGCCTCATCTA
CAACGTCAAGATCAGAGGGGTGAACTTCCCATCCAACGGCCCTGTGATGCAGAAGAAAACA
CTCGGCTGGGAGGCCAACACCGAGATGCTGTACCCCGCTGACGGCGGCCTGGAAGGCAG
AACCGACATGGCCCTGAAGCTCGTGGGCGGGGGCCACCTGATCTGCAACTTCAAGACCAC
ATACAGATCCAAGAAACCCGCTAAGAACCTCAAGATGCCCGGCGTCTACTATGTGGACCAC
AGACTGGAAAGAATCAAGGAGGCCGACAAAGAGACCTACGTCGAGCAGCACGAGGTGGCT
GTGGCCAGATACTGCGACCTCCCTAGCAAACTGGGGCACAAACTTAATGGCATGGACGAGC
TGTACAAGTCCGGACTCAGATCTCGAGCTCAAGCTTCGAACGAATTCGGCATTGATCTGAG
CGGCCTGACCCTGCAGGAATTCAGCTCCCCGAAACGCCCTAAGCTGCTGCGTGAATATTAC
GATTGGCTGGTTGATAACAGCTTTACCCCATATCTGGTGGTGGATGCCACATACCTGGGCGT
GAACGTGCCCGTGGAGTATGTGAAAGACGGTCAGATCGTGCTGAATCTGTCTGCAAGTGTG
ACCGGCAACCTGCAACTGACAAATGATTTTATCCAGTTCAACGCCCAGTTTAAGGGCGTGTC
TCGTGAACTGTATATCCCGATGGGTGCCGCTCTGGCCATTTACGCTCGCGAGAACGGCGAT
GGTGTGATGTTCGAACCAGAAGAAATCTATGACGAGCTGAATATTGGTCAGATCTCGTACGC
CTCCCGGGGCGGTGGCTCATCTGGCGGAGGTGAGCTCCAGACAATTAAGTGTGTTGTTGT
GGGCGATGGTGCTTAA **---IRES Region---**
ATGGGGGTTGGTAAAACATGTCTCCTGATATCCTACACAACAAACAAATTTCCATCGGAGTAT
GTACCGACTGTTTTTGACAACTATGCAGTCACAGTTATGATTGGTGGAGAACCATATACTCTT
GGACTTTTTGATACTGCAGGGCTAGAGGATTATGACAGATTACGACCGCTGAGTTATCCACA
AACAGATGTATTTCTAGTCTGTTTTTCAGTGGTCTCTCCATCTTCATTTGAAAACGTGAAAGA
AAAGTGGGTGCCTGAGATAACTCACCACTGTCCAAAGACTCCTTTCTTGCTTGTTGGGACT
CAAATTGATCTCAGAGATGACCCCTCTACTATTGAGAAACTTGCCAAGAACAAACAGAAGCC

TATCACTCCAGAGACTGCTGAAAAGCTGGCCCGTGACCTGAAGGCTGTCAAGTATGTGGAG
TGTTCTGCACTTACACAGAGAGGTCTGAAGAATGTGTTTGATGAGGCTATCCTAGCTGCCCT
CGAGCCTCCGGAAACTCAACCCGGGGATCCCAATTGGGAGCTCGTGTACACGGCGCGCCT
GCAGGGAGGTGGCTCATCTGGCGGAGGTCAGATCTCGTACGCGTCCCGGGGCGAGTTTC
TGGCAACCACACTGGAACGGATCGAGAAAAATTTCGTGATTACTGATCCGAGACTGCCTGA
CAACCCAATCATTTTTGCGAGCGATTCCTTCCTGCAGCTGACAGAATATTCTCGGGAAGAGA
TCCTGGGGCGCAATGCCCGTTTTCTGCAGGGACCCGAGACAGACCGTGCCACTGTTCGGA
AAATCAGAGATGCTATTGACAACCAGACTGAAGTGACCGTTCAGCTGATCAATTATACCAAG
AGCGGCAAGAAGTTCTGGAACGTGTTCCACCTGCAGCCGATGCGCGATTATAAGGGCGAC
GTCCAGTACTTCATTGGCGTGCAGCTGGATGGCACCGAACGTCTTCATGGCGCCGCTGAG
CGTGAGGCGGTCTGCCTGATCAAAAAGACAGCCTTTCAGATTGCTGAGGCAGCGAACGAC
GAAAATTACTTTGGAAGCGGGAGTGGGAGCGTGAGCAAGGGCGAGGAGCTGTTCACCGG
GGTGGTGCCCATCCTGGTCGAGCTGGACGGCGACGTAAACGGCCACAAGTTCAGCGTGTC
CGGCGAGGGCGAGGGCGATGCCACCTACGGCAAGCTGACCCTGAAGCTCATCTGCACCA
CCGGCAAGCTGCCCGTGCCCTGGCCCACCCTCGTGACCACCCTCGGCTACGGCCTGCAG
TGCTTCGCCCGCTACCCCGACCACATGAAGCAGCACGACTTCTTCAAGTCCGCCATGCCC
GAAGGCTACGTCCAGGAGCGCACCATCTTCTTCAAGGACGACGGCAACTACAAGACCCGC
GCCGAGGTGAAGTTCGAGGGCGACACCCTGGTGAACCGCATCGAGCTGAAGGGCATCGA
CTTCAAGGAGGACGGCAACATCCTGGGGCACAAGCTGGAGTACAACTACAACAGCCACAA
CGTCTATATCACCGCCGACAAGCAGAAGAACGGCATCAAGGCCAACTTCAAGATCCGCCAC
AACATCGAGGACGGCGGCGTGCAGCTCGCCGACCACTACCAGCAGAACACCCCCATCGG
CGACGGCCCCGTGCTGCTGCCCGACAACCACTACCTGAGCTACCAGTCCAAGCTGAGCAA
AGACCCCAACGAGAAGCGCGATCACATGGTCCTGCTGGAGTTCGTGACCGCCGCCGGGAT
CACTCTCGGCATGGACGAGCTGTACAAGAAAAAAAAGAAGAAAAAGAGCAAGACCAAATGC
GTGATTATGTAA

#### 7) pIRES-tagRFPt-SspB Micro-Rac1/N12-Rac1/13C-iLID-mVenus-CAAX

Amino Acid Sequence:

MVSKGEELIKENMHMKLYMEGTVNNHHFKCTSEGEGKPYEGTQTMRIKVVEGGPLPFAFDILA
TSFMYGSRTFINHTQGIPDFFKQSFPEGFTWERVTTYEDGGVLTATQDTSLQDGCLIYNVKIRG
VNFPSNGPVMQKKTLGWEANTEMLYPADGGLEGRTDMALKLVGGGHLICNFKTTYRSKKPAK
NLKMPGVYYVDHRLERIKEADKETYVEQHEVAVARYCDLPSKLGHKLNGMDELYKSGLRSRAQ
ASNEFGIDLSGLTLQEFSSPKRPKLLREYYDWLVDNSFTPYLVVDATYLGVNVPVEYVKDGQIVL
NLSASATGNLQLTNDFIQFNAQFKGVSRELYIPMGAALAIYARENGDGVMFEPEEIYDELNIGQIS
YASRGGGSSGGGELQAIKCVVVGDGA* **--- IRES Region---**
MVGKTCLLISYTTNAFPGEYIPTVFDNYSANVMVDGKPVNLGLWDTAGLEDYDRLRPLSYPQTD
VFLICFSLVSPASFENVRAKWYPEVRHHCPNTPIILVGTKLDLRDDKDTIEKLKEKKLTPITYPQGL
AMAKEIGAVKYLECSALTQRGLKTVFDEAIRAVLCPPPVGDPNWELVYTARLQGGGSSGGGQIS
YASRGEFLATTLERIEKNFVITDPRLPDNPIIFASDSFLQLTEYSREEILGRNCRFLQGPETDRATV
RKIRDAIDNQTEVTVQLINYTKSGKKFWNVFHLQPMRDYKGDVQYFIGVQLDGTERLHGAAER
EAVCLIKKTAFQIAEAANDENYFGSGSGSVSKGEELFTGVVPILVELDGDVNGHKFSVSGEGEG
DATYGKLTLKLICTTGKLPVPWPTLVTTLGYGLQCFARYPDHMKQHDFFKSAMPEGYVQERTIF
FKDDGNYKTRAEVKFEGDTLVNRIELKGIDFKEDGNILGHKLEYNYNSHNVYITADKQKNGIKAN
FKIRHNIEDGGVQLADHYQQNTPIGDGPVLLPDNHYLSYQSKLSKDPNEKRDHMVLLEFVTAAG
ITLGMDELYKKKKKKKSKTKCVIM*

DNA Sequence:

ATGGTGTCTAAGGGCGAAGAGCTGATTAAGGAGAACATGCACATGAAGCTGTACATGGAGG
GCACCGTGAACAACCACCACTTCAAGTGCACATCCGAGGGCGAAGGCAAGCCCTACGAGG
GCACCCAGACCATGAGAATCAAGGTGGTCGAGGGCGGCCCTCTCCCCTTCGCCTTCGACA
TCCTGGCTACCAGCTTCATGTACGGCAGCAGAACCTTCATCAACCACACCCAGGGCATCCC
CGACTTCTTTAAGCAGTCCTTCCCTGAGGGCTTCACATGGAGAGAGTCACCACATACGAA
GACGGGGGCGTGCTGACCGCTACCCAGGACACCAGCCTCCAGGACGGCTGCCTCATCTA
CAACGTCAAGATCAGAGGGGTGAACTTCCCATCCAACGGCCCTGTGATGCAGAAGAAAACA
CTCGGCTGGGAGGCCAACACCGAGATGCTGTACCCCGCTGACGGCGGCCTGGAAGGCAG
AACCGACATGGCCCTGAAGCTCGTGGGCGGGGGCCACCTGATCTGCAACTTCAAGACCAC
ATACAGATCCAAGAAACCCGCTAAGAACCTCAAGATGCCCGGCGTCTACTATGTGGACCAC
AGACTGGAAAGAATCAAGGAGGCCGACAAAGAGACCTACGTCGAGCAGCACGAGGTGGCT
GTGGCCAGATACTGCGACCTCCCTAGCAAACTGGGGCACAAACTTAATGGCATGGACGAGC
TGTACAAGTCCGGACTCAGATCTCGAGCTCAAGCTTCGAACGAATTCGGCATTGATCTGAG
CGGCCTGACCCTGCAGGAATTCAGCTCCCCGAAACGCCCTAAGCTGCTGCGTGAATATTAC
GATTGGCTGGTTGATAACAGCTTTACCCCATATCTGGTGGTGGATGCCACATACCTGGGCGT
GAACGTGCCCGTGGAGTATGTGAAAGACGGTCAGATCGTGCTGAATCTGTCTGCAAGTGC
GACCGGCAACCTGCAACTGACAAATGATTTTATCCAGTTCAACGCCCAGTTTAAGGGCGTG
TCTCGTGAACTGTATATCCCGATGGGTGCCGCTCTGGCCATTTACGCTCGCGAGAACGGCG
ATGGTGTGATGTTCGAACCAGAAGAAATCTATGACGAGCTGAATATTGGTCAGATCTCGTAC
GCCTCCCGGGGCGGTGGCTCATCTGGCGGAGGTGAGCTCCAGGCCATCAAGTGTGTGGT
GGTGGGAGACGGAGCTTAA **---IRES Region---**
ATGGTAGGTAAAACTTGCCTACTGATCAGTTACACAACCAATGCATTTCCTGGAGAATATATC
CCTACTGTCTTTGACAATTATTCTGCCAATGTTATGGTAGATGGAAAACCGGTGAATCTGGGC
TTATGGGATACAGCTGGACTAGAAGATTATGACAGATTACGCCCCCTATCCTATCCGCAAACA
GATGTGTTCTTAATTTGCTTTTCCCTTGTGAGTCCTGCATCATTTGAAAATGTCCGTGCAAAG
TGGTATCCTGAGGTGCGGCACCACTGTCCCAACACTCCCATCATCCTAGTGGGAACTAAAC
TTGATCTTAGGGATGATAAAGACACGATCGAGAAACTGAAGGAGAAGAAGCTGACTCCCATC

ACCTATCCGCAGGGTCTAGCCATGGCTAAGGAGATTGGTGCTGTAAAATACCTGGAGTGCT
CGGCGCTCACACAGCGAGGCCTCAAGACAGTGTTTGACGAAGCGATCCGAGCAGTCCTCT
GCCCGCCTCCCGTGGGGGATCCCAATTGGGAGCTCGTGTACACGGCGCGCCTGCAGGGA
GGTGGCTCATCTGGCGGAGGTCAGATCTCGTACGCGTCCCGGGGCGAGTTTCTGGCAACC
ACACTGGAACGGATCGAGAAAAATTTCGTGATTACTGATCCGAGACTGCCTGACAACCCAAT
CATTTTTGCGAGCGATTCCTTCCTGCAGCTGACAGAATATTCTCGGGAAGAGATCCTGGGG
CGCAATTGCCGTTTTCTGCAGGGACCCGAGACAGACCGTGCCACTGTTCGGAAAATCAGA
GATGCTATTGACAACCAGACTGAAGTGACCGTTCAGCTGATCAATTATACCAAGAGCGGCAA
GAAGTTCTGGAACGTGTTCCACCTGCAGCCGATGCGCGATTATAAGGGCGACGTCCAGTAC
TTCATTGGCGTGCAGCTGGATGGCACCGAACGTCTTCATGGCGCCGCTGAGCGTGAGGCG
GTCTGCCTGATCAAAAAGACAGCCTTTCAGATTGCTGAGGCAGCGAACGACGAAAATTACT
TTGGAAGCGGGAGTGGGAGCGTGAGCAAGGGCGAGGAGCTGTTCACCGGGGTGGTGCC
CATCCTGGTCGAGCTGGACGGCGACGTAAACGGCCACAAGTTCAGCGTGTCCGGCGAGG
GCGAGGGCGATGCCACCTACGGCAAGCTGACCCTGAAGCTCATCTGCACCACCGGCAAGC
TGCCCGTGCCCTGGCCCACCCTCGTGACCACCCTCGGCTACGGCCTGCAGTGCTTCGCC
CGCTACCCCGACCACATGAAGCAGCACGACTTCTTCAAGTCCGCCATGCCCGAAGGCTAC
GTCCAGGAGCGCACCATCTTCTTCAAGGACGACGGCAACTACAAGACCCGCGCCGAGGTG
AAGTTCGAGGGCGACACCCTGGTGAACCGCATCGAGCTGAAGGGCATCGACTTCAAGGAG
GACGGCAACATCCTGGGGCACAAGCTGGAGTACAACTACAACAGCCACAACGTCTATATCA
CCGCCGACAAGCAGAAGAACGGCATCAAGGCCAACTTCAAGATCCGCCACAACATCGAGG
ACGGCGGCGTGCAGCTCGCCGACCACTACCAGCAGAACACCCCCATCGGCGACGGCCCC
GTGCTGCTGCCCGACAACCACTACCTGAGCTACCAGTCCAAGCTGAGCAAAGACCCCAAC
GAGAAGCGCGATCACATGGTCCTGCTGGAGTTCGTGACCGCCGCCGGGATCACTCTCGGC
ATGGACGAGCTGTACAAGAAAAAAAAGAAGAAAAAGAGCAAGACCAAATGCGTGATTATGTA
A

#### 8) pIRES-tagRFPt-SspB Micro-Rac1/N12-Rac1/13C-iLID C252A-mVenus-CAAX

Amino Acid Sequence (C252A is underlined):

MVSKGEELIKENMHMKLYMEGTVNNHHFKCTSEGEGKPYEGTQTMRIKVVEGGPLPFAFDILA
TSFMYGSRTFINHTQGIPDFFKQSFPEGFTWERVTTYEDGGVLTATQDTSLQDGCLIYNVKIRG
VNFPSNGPVMQKKTLGWEANTEMLYPADGGLEGRTDMALKLVGGGHLICNFKTTYRSKKPAK
NLKMPGVYYVDHRLERIKEADKETYVEQHEVAVARYCDLPSKLGHKLNGMDELYKSGLRSRAQ
ASNEFGIDLSGLTLQEFSSPKRPKLLREYYDWLVDNSFTPYLVVDATYLGVNVPVEYVKDGQIVL
NLSASATGNLQLTNDFIQFNAQFKGVSRELYIPMGAALAIYARENGDGVMFEPEEIYDELNIGQIS
YASRGGGSSGGGELQAIKCVVVGDGA* **--- IRES Region---**
MVGKTCLLISYTTNAFPGEYIPTVFDNYSANVMVDGKPVNLGLWDTAGLEDYDRLRPLSYPQTD
VFLICFSLVSPASFENVRAKWYPEVRHHCPNTPIILVGTKLDLRDDKDTIEKLKEKKLTPITYPQGL
AMAKEIGAVKYLECSALTQRGLKTVFDEAIRAVLCPPPVGDPNWELVYTARLQGGGSSGGGQIS
YASRGEFLATTLERIEKNFVITDPRLPDNPIIFASDSFLQLTEYSREEILGRN<u>**A**</u>RFLQGPETDRATV
RKIRDAIDNQTEVTVQLINYTKSGKKFWNVFHLQPMRDYKGDVQYFIGVQLDGTERLHGAAER
EAVCLIKKTAFQIAEAANDENYFGSGSGSVSKGEELFTGVVPILVELDGDVNGHKFSVSGEGEG
DATYGKLTLKLICTTGKLPVPWPTLVTTLGYGLQCFARYPDHMKQHDFFKSAMPEGYVQERTIF
FKDDGNYKTRAEVKFEGDTLVNRIELKGIDFKEDGNILGHKLEYNYNSHNVYITADKQKNGIKAN
FKIRHNIEDGGVQLADHYQQNTPIGDGPVLLPDNHYLSYQSKLSKDPNEKRDHMVLLEFVTAAG
ITLGMDELYKKKKKKKSKTKCVIM*

DNA Sequence:

ATGGTGTCTAAGGGCGAAGAGCTGATTAAGGAGAACATGCACATGAAGCTGTACATGGAGG
GCACCGTGAACAACCACCACTTCAAGTGCACATCCGAGGGCGAAGGCAAGCCCTACGAGG
GCACCCAGACCATGAGAATCAAGGTGGTCGAGGGCGGCCCTCTCCCCTTCGCCTTCGACA
TCCTGGCTACCAGCTTCATGTACGGCAGCAGAACCTTCATCAACCACACCCAGGGCATCCC
CGACTTCTTTAAGCAGTCCTTCCCTGAGGGCTTCACATGGGAGAGAGTCACCACATACGAA
GACGGGGGCGTGCTGACCGCTACCCAGGACACCAGCCTCCAGGACGGCTGCCTCATCTA
CAACGTCAAGATCAGAGGGGTGAACTTCCCATCCAACGGCCCTGTGATGCAGAAGAAAACA
CTCGGCTGGGAGGCCAACACCGAGATGCTGTACCCCGCTGACGGCGGCCTGGAAGGCAG
AACCGACATGGCCCTGAAGCTCGTGGGCGGGGGCCACCTGATCTGCAACTTCAAGACCAC
ATACAGATCCAAGAAACCCGCTAAGAACCTCAAGATGCCCGGCGTCTACTATGTGGACCAC
AGACTGGAAAGAATCAAGGAGGCCGACAAAGAGACCTACGTCGAGCAGCACGAGGTGGCT
GTGGCCAGATACTGCGACCTCCCTAGCAAACTGGGGCACAAACTTAATGGCATGGACGAGC
TGTACAAGTCCGGACTCAGATCTCGAGCTCAAGCTTCGAACGAATTCGGCATTGATCTGAG
CGGCCTGACCCTGCAGGAATTCAGCTCCCCGAAACGCCCTAAGCTGCTGCGTGAATATTAC
GATTGGCTGGTTGATAACAGCTTTACCCCATATCTGGTGGTGGATGCCACATACCTGGGCGT
GAACGTGCCCGTGGAGTATGTGAAAGACGGTCAGATCGTGCTGAATCTGTCTGCAAGTGC
GACCGGCAACCTGCAACTGACAAATGATTTTATCCAGTTCAACGCCCAGTTTAAGGGCGTG
TCTCGTGAACTGTATATCCCGATGGGTGCCGCTCTGGCCATTTACGCTCGCGAGAACGGCG
ATGGTGTGATGTTCGAACCAGAAGAAATCTATGACGAGCTGAATATTGGTCAGATCTCGTAC
GCCTCCCGGGGCGGTGGCTCATCTGGCGGAGGTGAGCTCCAGGCCATCAAGTGTGTGGT
GGTGGGAGACGGAGCTTAA **---IRES Region---**
ATGGTAGGTAAAACTTGCCTACTGATCAGTTACACAACCAATGCATTTCCTGGAGAATATATC
CCTACTGTCTTTGACAATTATTCTGCCAATGTTATGGTAGATGGAAAACCGGTGAATCTGGGC
TTATGGGATACAGCTGGACTAGAAGATTATGACAGATTACGCCCCCTATCCTATCCGCAAACA
GATGTGTTCTTAATTTGCTTTTCCCTTGTGAGTCCTGCATCATTTGAAAATGTCCGTGCAAAG
TGGTATCCTGAGGTGCGGCACCACTGTCCCAACACTCCCATCATCCTAGTGGGAACTAAAC
TTGATCTTAGGGATGATAAAGACACGATCGAGAAACTGAAGGAGAAGAAGCTGACTCCCATC

ACCTATCCGCAGGGTCTAGCCATGGCTAAGGAGATTGGTGCTGTAAAATACCTGGAGTGCT
CGGCGCTCACACAGCGAGGCCTCAAGACAGTGTTTGACGAAGCGATCCGAGCAGTCCTCT
GCCCGCCTCCCGTGGGGGATCCCAATTGGGAGCTCGTGTACACGGCGCGCCTGCAGGGA
GGTGGCTCATCTGGCGGAGGTCAGATCTCGTACGCGTCCCGGGGCGAGTTTCTGGCAACC
ACACTGGAACGGATCGAGAAAAATTTCGTGATTACTGATCCGAGACTGCCTGACAACCCAAT
CATTTTTGCGAGCGATTCCTTCCTGCAGCTGACAGAATATTCTCGGGAAGAGATCCTGGGG
CGCAATGCCCGTTTTCTGCAGGGACCCGAGACAGACCGTGCCACTGTTCGGAAAATCAGA
GATGCTATTGACAACCAGACTGAAGTGACCGTTCAGCTGATCAATTATACCAAGAGCGGCAA
GAAGTTCTGGAACGTGTTCCACCTGCAGCCGATGCGCGATTATAAGGGCGACGTCCAGTAC
TTCATTGGCGTGCAGCTGGATGGCACCGAACGTCTTCATGGCGCCGCTGAGCGTGAGGCG
GTCTGCCTGATCAAAAAGACAGCCTTTCAGATTGCTGAGGCAGCGAACGACGAAAATTACT
TTGGAAGCGGGAGTGGGAGCGTGAGCAAGGGCGAGGAGCTGTTCACCGGGGTGGTGCC
CATCCTGGTCGAGCTGGACGGCGACGTAAACGGCCACAAGTTCAGCGTGTCCGGCGAGG
GCGAGGGCGATGCCACCTACGGCAAGCTGACCCTGAAGCTCATCTGCACCACCGGCAAGC
TGCCCGTGCCCTGGCCCACCCTCGTGACCACCCTCGGCTACGGCCTGCAGTGCTTCGCC
CGCTACCCCGACCACATGAAGCAGCACGACTTCTTCAAGTCCGCCATGCCCGAAGGCTAC
GTCCAGGAGCGCACCATCTTCTTCAAGGACGACGGCAACTACAAGACCCGCGCCGAGGTG
AAGTTCGAGGGCGACACCCTGGTGAACCGCATCGAGCTGAAGGGCATCGACTTCAAGGAG
GACGGCAACATCCTGGGGCACAAGCTGGAGTACAACTACAACAGCCACAACGTCTATATCA
CCGCCGACAAGCAGAAGAACGGCATCAAGGCCAACTTCAAGATCCGCCACAACATCGAGG
ACGGCGGCGTGCAGCTCGCCGACCACTACCAGCAGAACACCCCCATCGGCGACGGCCCC
GTGCTGCTGCCCGACAACCACTACCTGAGCTACCAGTCCAAGCTGAGCAAAGACCCCAAC
GAGAAGCGCGATCACATGGTCCTGCTGGAGTTCGTGACCGCCGCCGGGATCACTCTCGGC
ATGGACGAGCTGTACAAGAAAAAAAAGAAGAAAAAGAGCAAGACCAAATGCGTGATTATGTA
A

#### 9) pIRES-tagRFPt-SspB Micro-RhoA/N14-RhoA/15C-iLID-mVenus-CAAX

Amino Acid Sequence:

MVSKGEELIKENMHMKLYMEGTVNNHHFKCTSEGEGKPYEGTQTMRIKVVEGGPLPFAFDILA
TSFMYGSRTFINHTQGIPDFFKQSFPEGFTWERVTTYEDGGVLTATQDTSLQDGCLIYNVKIRG
VNFPSNGPVMQKKTLGWEANTEMLYPADGGLEGRTDMALKLVGGGHLICNFKTTYRSKKPAK
NLKMPGVYYVDHRLERIKEADKETYVEQHEVAVARYCDLPSKLGHKLNGMDELYKSGLRSRAQ
ASNEFGIDLSGLTLQEFSSPKRPKLLREYYDWLVDNSFTPYLVVDATYLGVNVPVEYVKDGQIVL
NLSASATGNLQLTNDFIQFNAQFKGVSRELYIPMGAALAIYARENGDGVMFEPEEIYDELNIGQIS
YASRGGGSSGGGELAAIRKKLVIVGDGA* **---IRES Region---**
MCGKTCLLIVFSKDQFPEVYVPTVFENYVADIEVDGKQVELALWDTAGLEDYDRLRPLSYPDTD
VILMCFSIDSPDSLENIPEKWTPEVKHFCPNVPIILVGNKKDLRNDEHTRRELAKMKQEPVKPEE
GRDMANRIGAFGYMECSAKTKDGVREVFEMATRAALQAGDPNWELVYTARLQGGGSSGGGQ
ISYASRGEFLATTLERIEKNFVITDPRLPDNPIIFASDSFLQLTEYSREEILGRNCRFLQGPETDRA
TVRKIRDAIDNQTEVTVQLINYTKSGKKFWNVFHLQPMRDYKGDVQYFIGVQLDGTERLHGAA
EREAVCLIKKTAFQIAEAANDENYFGSGSGSVSKGEELFTGVVPILVELDGDVNGHKFSVSGEG
EGDATYGKLTLKLICTTGKLPVPWPTLVTTLGYGLQCFARYPDHMKQHDFFKSAMPEGYVQER
TIFFKDDGNYKTRAEVKFEGDTLVNRIELKGIDFKEDGNILGHKLEYNYNSHNVYITADKQKNGIK
ANFKIRHNIEDGGVQLADHYQQNTPIGDGPVLLPDNHYLSYQSKLSKDPNEKRDHMVLLEFVTA
AGITLGMDELYKKKKKKKSKTKCVIM*

DNA Sequence:

ATGGTGTCTAAGGGCGAAGAGCTGATTAAGGAGAACATGCACATGAAGCTGTACATGGAGG
GCACCGTGAACAACCACCACTTCAAGTGCACATCCGAGGGCGAAGGCAAGCCCTACGAGG
GCACCCAGACCATGAGAATCAAGGTGGTCGAGGGCGGCCCTCTCCCCTTCGCCTTCGACA
TCCTGGCTACCAGCTTCATGTACGGCAGCAGAACCTTCATCAACCACACCCAGGGCATCCC
CGACTTCTTTAAGCAGTCCTTCCCTGAGGGCTTCACATGGGAGAGAGTCACCACATACGAA
GACGGGGGCGTGCTGACCGCTACCCAGGACACCAGCCTCCAGGACGGCTGCCTCATCTA
CAACGTCAAGATCAGAGGGGTGAACTTCCCATCCAACGGCCCTGTGATGCAGAAGAAAACA
CTCGGCTGGGAGGCCAACACCGAGATGCTGTACCCCGCTGACGGCGGCCTGGAAGGCAG
AACCGACATGGCCCTGAAGCTCGTGGGCGGGGGCCACCTGATCTGCAACTTCAAGACCAC
ATACAGATCCAAGAAACCCGCTAAGAACCTCAAGATGCCCGGCGTCTACTATGTGGACCAC
AGACTGGAAAGAATCAAGGAGGCCGACAAAGAGACCTACGTCGAGCAGCACGAGGTGGCT
GTGGCCAGATACTGCGACCTCCCTAGCAAACTGGGGCACAAACTTAATGGCATGGACGAGC
TGTACAAGTCCGGACTCAGATCTCGAGCTCAAGCTTCGAACGAATTCGGCATTGATCTGAG
CGGCCTGACCCTGCAGGAATTCAGCTCCCCGAAACGCCCTAAGCTGCTGCGTGAATATTAC
GATTGGCTGGTTGATAACAGCTTTACCCCATATCTGGTGGTGGATGCCACATACCTGGGCGT
GAACGTGCCCGTGGAGTATGTGAAAGACGGTCAGATCGTGCTGAATCTGTCTGCAAGTGC
GACCGGCAACCTGCAACTGACAAATGATTTTATCCAGTTCAACGCCCAGTTTAAGGGCGTG
TCTCGTGAACTGTATATCCCGATGGGTGCCGCTCTGGCCATTTACGCTCGCGAGAACGGCG
ATGGTGTGATGTTCGAACCAGAAGAAATCTATGACGAGCTGAATATTGGTCAGATCTCGTAC
GCCTCCCGGGGCGGTGGCTCATCTGGCGGAGGTGAGCTCGCTGCCATCCGGAAGAAACT
GGTGATTGTTGGTGATGGAGCCTAA **---IRES Region---**
ATGTGTGGAAAGACATGCTTGCTCATAGTCTTCAGCAAGGATCAGTTCCCAGAGGTGTATGT
GCCCACAGTGTTTGAGAACTATGTGGCAGATATCGAGGTGGATGGAAAGCAGGTAGAGTTG
GCTTTGTGGGACACAGCTGGGCTGGAAGATTATGATCGCCTGAGGCCCCTCTCCTACCCAG
ATACCGATGTTATACTGATGTGTTTTTCCATCGACAGCCCTGATAGTTTAGAAAACATCCCAG
AAAAGTGGACCCCAGAAGTCAAGCATTTCTGTCCCAACGTGCCCATCATCCTGGTTGGGAA
TAAGAAGGATCTTCGGAATGATGAGCACACAAGGCGGGAGCTAGCCAAGATGAAGCAGGA

GCCGGTGAAACCTGAAGAAGGCAGAGATATGGCAAACAGGATTGGCGCTTTTGGGTACATG
GAGTGTTCAGCAAAGACCAAAGATGGAGTGAGAGAGGTTTTTGAAATGGCTACGAGAGCTG
CTCTGCAAGCTGGGGATCCCAATTGGGAGCTCGTGTACACGGCGCGCCTGCAGGGAGGT
GGCTCATCTGGCGGAGGTCAGATCTCGTACGCGTCCCGGGGCGAGTTTCTGGCAACCACA
CTGGAACGGATCGAGAAAAATTTCGTGATTACTGATCCGAGACTGCCTGACAACCCAATCAT
TTTTGCGAGCGATTCCTTCCTGCAGCTGACAGAATATTCTCGGGAAGAGATCCTGGGGCGC
AATTGCCGTTTTCTGCAGGGACCCGAGACAGACCGTGCCACTGTTCGGAAAATCAGAGATG
CTATTGACAACCAGACTGAAGTGACCGTTCAGCTGATCAATTATACCAAGAGCGGCAAGAAG
TTCTGGAACGTGTTCCACCTGCAGCCGATGCGCGATTATAAGGGCGACGTCCAGTACTTCA
TTGGCGTGCAGCTGGATGGCACCGAACGTCTTCATGGCGCCGCTGAGCGTGAGGCGGTC
TGCCTGATCAAAAAGACAGCCTTTCAGATTGCTGAGGCAGCGAACGACGAAAATTACTTTG
GAAGCGGGAGTGGGAGCGTGAGCAAGGGCGAGGAGCTGTTCACCGGGGTGGTGCCCAT
CCTGGTCGAGCTGGACGGCGACGTAAACGGCCACAAGTTCAGCGTGTCCGGCGAGGGCG
AGGGCGATGCCACCTACGGCAAGCTGACCCTGAAGCTCATCTGCACCACCGGCAAGCTGC
CCGTGCCCTGGCCCACCCTCGTGACCACCCTCGGCTACGGCCTGCAGTGCTTCGCCCGC
TACCCCGACCACATGAAGCAGCACGACTTCTTCAAGTCCGCCATGCCCGAAGGCTACGTCC
AGGAGCGCACCATCTTCTTCAAGGACGACGGCAACTACAAGACCCGCGCCGAGGTGAAGT
TCGAGGGCGACACCCTGGTGAACCGCATCGAGCTGAAGGGCATCGACTTCAAGGAGGAC
GGCAACATCCTGGGGCACAAGCTGGAGTACAACTACAACAGCCACAACGTCTATATCACCG
CCGACAAGCAGAAGAACGGCATCAAGGCCAACTTCAAGATCCGCCACAACATCGAGGACG
GCGGCGTGCAGCTCGCCGACCACTACCAGCAGAACACCCCCATCGGCGACGGCCCCGTG
CTGCTGCCCGACAACCACTACCTGAGCTACCAGTCCAAGCTGAGCAAAGACCCCAACGAG
AAGCGCGATCACATGGTCCTGCTGGAGTTCGTGACCGCCGCCGGGATCACTCTCGGCATG
GACGAGCTGTACAAGAAAAAAAAGAAGAAAAAGAGCAAGACCAAATGCGTGATTATGTAA

**10) pIRES-tagRFPt-SspB Micro-RhoA/N14-RhoA/15C-iLID C252A-mVenus-CAAX**

Amino Acid Sequence (C252A is underlined):

MVSKGEELIKENMHMKLYMEGTVNNHHFKCTSEGEGKPYEGTQTMRIKVVEGGPLPFAFDILA
TSFMYGSRTFINHTQGIPDFFKQSFPEGFTWERVTTYEDGGVLTATQDTSLQDGCLIYNVKIRG
VNFPSNGPVMQKKTLGWEANTEMLYPADGGLEGRTDMALKLVGGGHLICNFKTTYRSKKPAK
NLKMPGVYYVDHRLERIKEADKETYVEQHEVAVARYCDLPSKLGHKLNGMDELYKSGLRSRAQ
ASNEFGIDLSGLTLQEFSSPKRPKLLREYYDWLVDNSFTPYLVVDATYLGVNVPVEYVKDGQIVL
NLSASATGNLQLTNDFIQFNAQFKGVSRELYIPMGAALAIYARENGDGVMFEPEEIYDELNIGQIS
YASRGGGSSGGGELAAIRKKLVIVGDGA* **---IRES Region---**
MCGKTCLLIVFSKDQFPEVYVPTVFENYVADIEVDGKQVELALWDTAGLEDYDRLRPLSYPDTD
VILMCFSIDSPDSLENIPEKWTPEVKHFCPNVPIILVGNKKDLRNDEHTRRELAKMKQEPVKPEE
GRDMANRIGAFGYMECSAKTKDGVREVFEMATRAALQAGDPNWELVYTARLQGGGSSGGGQ
ISYASRGEFLATTLERIEKNFVITDPRLPDNPIIFASDSFLQLTEYSREEILGRN<u>**A**</u>RFLQGPETDRA
TVRKIRDAIDNQTEVTVQLINYTKSGKKFWNVFHLQPMRDYKGDVQYFIGVQLDGTERLHGAA
EREAVCLIKKTAFQIAEAANDENYFGSGSGSVSKGEELFTGVVPILVELDGDVNGHKFSVSGEG
EGDATYGKLTLKLICTTGKLPVPWPTLVTTLGYGLQCFARYPDHMKQHDFFKSAMPEGYVQER
TIFFKDDGNYKTRAEVKFEGDTLVNRIELKGIDFKEDGNILGHKLEYNYNSHNVYITADKQKNGIK
ANFKIRHNIEDGGVQLADHYQQNTPIGDGPVLLPDNHYLSYQSKLSKDPNEKRDHMVLLEFVTA
AGITLGMDELYKKKKKKKSKTKCVIM*

DNA Sequence:

ATGGTGTCTAAGGGCGAAGAGCTGATTAAGGAGAACATGCACATGAAGCTGTACATGGAGG
GCACCGTGAACAACCACCACTTCAAGTGCACATCCGAGGGCGAAGGCAAGCCCTACGAGG
GCACCCAGACCATGAGAATCAAGGTGGTCGAGGGCGGCCCTCTCCCCTTCGCCTTCGACA
TCCTGGCTACCAGCTTCATGTACGGCAGCAGAACCTTCATCAACCACACCCAGGGCATCCC
CGACTTCTTTAAGCAGTCCTTCCCTGAGGGCTTCACATGGGAGAGAGTCACCACATACGAA
GACGGGGGCGTGCTGACCGCTACCCAGGACACCAGCCTCCAGGACGGCTGCCTCATCTA
CAACGTCAAGATCAGAGGGGTGAACTTCCCATCCAACGGCCCTGTGATGCAGAAGAAAACA
CTCGGCTGGGAGGCCAACACCGAGATGCTGTACCCCGCTGACGGCGGCCTGGAAGGCAG
AACCGACATGGCCCTGAAGCTCGTGGGCGGGGGCCACCTGATCTGCAACTTCAAGACCAC
ATACAGATCCAAGAAACCCGCTAAGAACCTCAAGATGCCCGGCGTCTACTATGTGGACCAC
AGACTGGAAAGAATCAAGGAGGCCGACAAAGAGACCTACGTCGAGCAGCACGAGGTGGCT
GTGGCCAGATACTGCGACCTCCCTAGCAAACTGGGGCACAAACTTAATGGCATGGACGAGC
TGTACAAGTCCGGACTCAGATCTCGAGCTCAAGCTTCGAACGAATTCGGCATTGATCTGAG
CGGCCTGACCCTGCAGGAATTCAGCTCCCCGAAACGCCCTAAGCTGCTGCGTGAATATTAC
GATTGGCTGGTTGATAACAGCTTTACCCCATATCTGGTGGTGGATGCCACATACCTGGGCGT
GAACGTGCCCGTGGAGTATGTGAAAGACGGTCAGATCGTGCTGAATCTGTCTGCAAGTGC
GACCGGCAACCTGCAACTGACAAATGATTTTATCCAGTTCAACGCCCAGTTTAAGGGCGTG
TCTCGTGAACTGTATATCCCGATGGGTGCCGCTCTGGCCATTTACGCTCGCGAGAACGGCG
ATGGTGTGATGTTCGAACCAGAAGAAATCTATGACGAGCTGAATATTGGTCAGATCTCGTAC
GCCTCCCGGGGCGGTGGCTCATCTGGCGGAGGTGAGCTCGCTGCCATCCGGAAGAAACT
GGTGATTGTTGGTGATGGAGCCTAA **---IRES Region---**
ATGTGTGGAAAGACATGCTTGCTCATAGTCTTCAGCAAGGATCAGTTCCCAGAGGTGTATGT
GCCCACAGTGTTTGAGAACTATGTGGCAGATATCGAGGTGGATGGAAAGCAGGTAGAGTTG
GCTTTGTGGGACACAGCTGGGCTGGAAGATTATGATCGCCTGAGGCCCCTCTCCTACCCAG
ATACCGATGTTATACTGATGTGTTTTTCCATCGACAGCCCTGATAGTTTAGAAAACATCCCAG
AAAAGTGGACCCCAGAAGTCAAGCATTTCTGTCCCAACGTGCCCATCATCCTGGTTGGGAA
TAAGAAGGATCTTCGGAATGATGAGCACACAAGGCGGGAGCTAGCCAAGATGAAGCAGGA

GCCGGTGAAACCTGAAGAAGGCAGAGATATGGCAAACAGGATTGGCGCTTTTGGGTACATG
GAGTGTTCAGCAAAGACCAAAGATGGAGTGAGAGAGGTTTTTGAAATGGCTACGAGAGCTG
CTCTGCAAGCTGGGGATCCCAATTGGGAGCTCGTGTACACGGCGCGCCTGCAGGGAGGT
GGCTCATCTGGCGGAGGTCAGATCTCGTACGCGTCCCGGGGCGAGTTTCTGGCAACCACA
CTGGAACGGATCGAGAAAAATTTCGTGATTACTGATCCGAGACTGCCTGACAACCCAATCAT
TTTTGCGAGCGATTCCTTCCTGCAGCTGACAGAATATTCTCGGGAAGAGATCCTGGGGCGC
AATGCCCGTTTTCTGCAGGGACCCGAGACAGACCGTGCCACTGTTCGGAAAATCAGAGAT
GCTATTGACAACCAGACTGAAGTGACCGTTCAGCTGATCAATTATACCAAGAGCGGCAAGAA
GTTCTGGAACGTGTTCCACCTGCAGCCGATGCGCGATTATAAGGGCGACGTCCAGTACTTC
ATTGGCGTGCAGCTGGATGGCACCGAACGTCTTCATGGCGCCGCTGAGCGTGAGGCGGT
CTGCCTGATCAAAAAGACAGCCTTTCAGATTGCTGAGGCAGCGAACGACGAAAATTACTTT
GGAAGCGGGAGTGGGAGCGTGAGCAAGGGCGAGGAGCTGTTCACCGGGGTGGTGCCCA
TCCTGGTCGAGCTGGACGGCGACGTAAACGGCCACAAGTTCAGCGTGTCCGGCGAGGGC
GAGGGCGATGCCACCTACGGCAAGCTGACCCTGAAGCTCATCTGCACCACCGGCAAGCTG
CCCGTGCCCTGGCCCACCCTCGTGACCACCCTCGGCTACGGCCTGCAGTGCTTCGCCCG
CTACCCCGACCACATGAAGCAGCACGACTTCTTCAAGTCCGCCATGCCCGAAGGCTACGTC
CAGGAGCGCACCATCTTCTTCAAGGACGACGGCAACTACAAGACCCGCGCCGAGGTGAAG
TTCGAGGGCGACACCCTGGTGAACCGCATCGAGCTGAAGGGCATCGACTTCAAGGAGGAC
GGCAACATCCTGGGGCACAAGCTGGAGTACAACTACAACAGCCACAACGTCTATATCACCG
CCGACAAGCAGAAGAACGGCATCAAGGCCAACTTCAAGATCCGCCACAACATCGAGGACG
GCGGCGTGCAGCTCGCCGACCACTACCAGCAGAACACCCCCATCGGCGACGGCCCCGTG
CTGCTGCCCGACAACCACTACCTGAGCTACCAGTCCAAGCTAAGCAAAGACCCCAACGAG
AAGCGCGATCACATGGTCCTGCTGGAGTTCGTGACCGCCGCCGGGATCACTCTCGGCATG
GACGAGCTGTACAAGAAAAAAAAGAAGAAAAAGAGCAAGACCAAATGCGTGATTATGTAA
